# *In vitro* and *in vivo* characterization of a recombinant rhesus cytomegalovirus containing a complete genome

**DOI:** 10.1101/2020.06.02.129486

**Authors:** Husam Taher, Eisa Mahyari, Craig Kreklywich, Luke S. Uebelhoer, Matthew R. McArdle, Matilda J. Moström, Amruta Bhusari, Michael Nekorchuk, Travis Whitmer, Elizabeth A. Scheef, Lesli M. Sprehe, Dawn Roberts, Colette M. Hughes, Kerianne A. Jackson, Andrea N. Selseth, Abigail B. Ventura, Yujuan Yue, Kimberli A. Schmidt, Jason Shao, Paul T. Edlefsen, Jeremy Smedley, Richard J. Stanton, Michael K. Axthelm, Jacob D. Estes, Scott G. Hansen, Amitinder Kaur, Peter A. Barry, Benjamin N. Bimber, Louis J. Picker, Daniel N. Streblow, Klaus Früh, Daniel Malouli

**Affiliations:** Vaccine and Gene Therapy Institute, Oregon Health and Science University, Beaverton, Oregon, United States of America; Oregon National Primate Research Center, Oregon Health and Science University, Beaverton, Oregon, United States of America; Tulane National Primate Research Center, Covington, LA, United States; Division of Infection and Immunity, Cardiff University School of Medicine, Cardiff, United Kingdom; Center for Comparative Medicine and Department of Medical Pathology, University of California, Davis, California, USA; Statistical Center for HIV/AIDS Research and Prevention, Vaccine and Infectious Disease Division, Fred Hutchinson Cancer Research Center, Seattle, Washington, USA

**Author notes:** These authors contributed equally (Husam Taher, Eisa Mahyari). Department of Pediatrics, Oregon Health & Science University, Portland, Oregon, USA. Department of Biochemistry, University of Utah, Salt Lake City, Utah, USA. Corresponding author: Daniel Malouli, Vaccine and Gene Therapy Institute, Oregon Health and Science University, 505 NW 185^th^ Ave., Beaverton, OR 97006.

## Abstract

Cytomegaloviruses (CMVs) are highly adapted to their host species resulting in strict species specificity. Hence, *in vivo* examination of all aspects of CMV biology employs animal models using host-specific CMVs. Infection of rhesus macaques (RM) with rhesus CMV (RhCMV) has been established as a representative model for infection of humans with HCMV due to the close evolutionary relationships of both host and virus. However, the commonly used 68-1 strain of RhCMV has been passaged in fibroblasts for decades resulting in multiple genomic changes due to tissue culture adaptation that cause reduced viremia in RhCMV-naïve animals and limited shedding compared to low passage isolates. Using sequence information from primary RhCMV isolates we constructed a full-length (FL) RhCMV by repairing all presumed mutations in the 68-1 bacterial artificial chromosome (BAC). Inoculation of adult, immunocompetent, RhCMV-naïve RM with the reconstituted virus resulted in significant replication in the blood similar to primary isolates of RhCMV and furthermore led to extensive viremia in many tissues at day 14 post infection. In contrast, viral dissemination and viremia was greatly reduced upon deletion of genes also lacking in 68-1. Transcriptome analysis of infected tissues further revealed that chemokine-like genes deleted in 68-1 are among the most highly expressed viral transcripts both *in vitro* and *in vivo* consistent with an important immunomodulatory function of the respective proteins. We conclude that FL-RhCMV displays *in vitro* and *in vivo* characteristics of a wildtype virus while being amenable to genetic modifications through BAC recombineering techniques.

**Author Summary:** Human cytomegalovirus (HCMV) infections are generally asymptomatic in healthy immunocompetent individuals, but HCMV can cause serious disease after congenital infection and in individuals with immunocompromised immune systems. Since HCMV is highly species specific and cannot productively infect immunocompetent laboratory animals, experimental infection of rhesus macaques (RM) with rhesus CMV (RhCMV) has been established as a closely related animal model for HCMV. By employing the unique ability of CMV to elicit robust and lasting cellular immunity, this model has also been instrumental in developing novel CMV-based vaccines against chronic and recurring infections with pathogens such as the human immunodeficiency virus (HIV) and *Mycobacterium tuberculosis (Mtb)*. However, most of this work was conducted with derivatives of the 68-1 strain of RhCMV which has acquired multiple genomic alterations in tissue culture. To model pathogenesis and immunology of clinical HCMV isolates we generated a full-length (FL) RhCMV clone representative of low passage isolates. Infection of RhCMV-naïve RM with FL-RhCMV demonstrated viremia and tissue dissemination that was comparable to that of non-clonal low passage isolates. We further demonstrate that FL-RhCMV is strongly attenuated upon deletion of gene regions absent in 68-1 thus demonstrating the usefulness of FL-RhCMV to study RhCMV pathogenesis.

## Introduction

Chronic human cytomegalovirus (HCMV) infections are generally asymptomatic in healthy, immunocompetent individuals and seroprevalence ranges from approximately 45% in developed countries to almost 100% of the population in the developing world (1). However, the virus can cause significant disease after congenital infection and in individuals with immunocompromised immune systems (2). No vaccines against HCMV exist, and treatment with antiviral drugs can limit acute infections but cannot eliminate the persistent virus (3). Cytomegaloviruses are double stranded DNA viruses belonging to the herpesvirus subfamily *Betaherpesvirinae* and have so far been exclusively found in mammals, mainly rodents and primates (4). CMVs contain the largest genomes of all herpesviruses and current annotations predict upwards of 170 open reading frames (ORFs) for most species. Ribozyme profiling data suggests that the actual number of translated viral mRNAs is likely significantly higher (5), however only a subset of these produce high levels of protein during infection of fibroblasts (6, 7). Co-evolution of these viruses with their host species over millions of years has led to a sequence relationship between CMV species that generally mirrors that of their hosts while also resulting in strict species specificity (8, 9). Hence, HCMV does not replicate and is not pathogenic in immunocompetent animals, and animal models of HCMV thus generally rely on studying infection of a given host with their respective animal CMV. The most commonly used models are mice, rats, guinea pigs and rhesus macaques (RM). The close evolutionary relationship of RM to humans (as compared to rodents) is mirrored in the evolutionary relationship of the rhesus CMV (RhCMV) genome to HCMV as the overall genomic organization is similar and most viral gene families are found in both CMV species (10).

Infection of RM with RhCMV has thus become a highly useful animal model for HCMV including a model for congenital infection (11). In addition, RhCMV has been used extensively to explore the possibility of harnessing the unique immune biology of CMV as a novel vaccine strategy, in particular the ability to elicit and maintain high frequencies of effector memory T cells (12). This work revealed not only that RhCMV-based vectors are remarkably effective in protecting RM against challenge with simian immunodeficiency virus (SIV), *Mtb* and *Plasmodium knowlesi* (13–16), but also uncovered a unique and unexpected ability of RhCMV to be genetically programmed to elicit CD8^+^ T cells that differ in their MHC restriction (17, 18). Importantly, highly attenuated RhCMV vaccine vectors that display highly reduced viremia, dissemination and shedding maintain the adaptive immune program and the ability to protect against pathogen challenge (19, 20).

However, the vast majority of these immunological and challenge studies relied on a molecular clone of RhCMV that was derived from strain 68-1 which differs significantly from circulating RhCMV strains. The RhCMV strain 68-1 was originally isolated in 1968 from the urine of a healthy RM (21) and had been extensively passaged on fibroblasts for more than 30 years before being cloned as a BAC (22). During this time, 68-1 has acquired multiple tissue culture adaptations (10) including an inversion in a genomic region homologous to the HCMV ULb’ region. This inversion simultaneously deleted the genes Rh157.5 and Rh157.4 (UL128 and UL130 in HCMV), two members of the viral pentameric receptor complex (PRC), as well as three of six genes encoding chemokine-like proteins homologous to the HCMV UL146 family (23). Similar to PRC-deficient HCMV, the loss of a functional PRC resulted in restricted cell tropism of 68-1 RhCMV *in vitro* (24, 25). PRC-dependent infection of non-fibroblast cells, such as epithelial and endothelial cells, was increased upon the insertion of the Rh157.5 and Rh157.4 genes obtained from the unrelated RhCMV 180.92 strain (25). Furthermore, strain 68-1 showed reduced viremia and shedding compared to the low passage isolates UCD52 and UCD59 upon primary infection of RhCMV-seronegative RM (26). UCD52, UCD59 and 180.92 have also been used in congenital infection studies (11, 27, 28). However, UCD52 and UCD59 (29, 30) represent non-clonal isolates that have been passaged on rhesus epithelial cells instead of fibroblasts, a culture methods that preserves the PRC but can also lead to tissue culture adaptations (31, 32). The 180.92 strain was shown to consist of a mixture of a tissue culture adapted and wildtype variants with the latter rapidly emerging as the dominant variant *in vivo* (33). Thus, there is a need for the construction of a BAC-cloned RhCMV representative of primary isolates to enable studies that reflect circulating RhCMV strains and recapitulate the pathogenesis of HCMV. In addition, such a tissue culture non-adapted, but genetically modifiable RhCMV clone would also be a useful tool to model HCMV-based vaccine development for live-attenuated candidates derived from clinical isolates (34).

Here, we describe the construction of such a BAC-cloned RhCMV genome in which all presumed mutations in 68-1 that result in altered ORFs were repaired thus closely reflecting a clone of the original 68-1 isolate prior to tissue culture passage. We demonstrate that the resulting viral sequence, termed FL-RhCMV, is representative of contemporary RhCMV isolates from multiple primate centers. FL-RhCMV demonstrates *in vitro* growth characteristics that are very similar to those reported for primary isolates of HCMV, including the rapid accumulation of mutations in the gene homologous to HCMV RL13. Furthermore, we show that FL-RhCMV displays wildtype-like viremia in RhCMV-seronegative RM. The availability of the first RhCMV BAC clone containing a complete genome sequence granting the derived virus all characteristics of a circulating isolate will enable the selected modulation of tissue tropism, pathogenesis and immune stimulation. This is exemplified by our demonstration that the deletion of the RhCMV homologs of HCMV UL128, UL130 and UL146 profoundly impacted viral dissemination and proliferation during acute infection. Thus, we report the generation of a RhCMV BAC that represents a primary isolate and that can serve as a modifiable progenitor for studies using RhCMV as model for HCMV infection or HCMV-based vaccine vectors.

## Results

### Construction of a full length (FL) RhCMV BAC and *in vitro* characterization

Compared to circulating and low passage isolates, RhCMV strain 68-1 has acquired a large inversion in the region homologous to one end of the HCMV “unique long” (U_L_) sequence of the genome (commonly referred to as the ULb’ region), flanked by deletions of multiple ORFs on either side of the inversion (23, 35, 36). When sequencing the clonal bacterial artificial chromosome (BAC) of 68-1, we additionally identified multiple viral ORFs that contained point mutations predicted to result in frameshifts or premature terminations of the annotated proteins (10). One of these point mutations is located in Rh61/Rh60 (UL36 in HCMV) and renders the encoded inhibitor of extrinsic apoptosis non-functional (37). By reversing the frameshift in Rh61/Rh60 and by inserting the missing PRC members Rh157.5 and Rh157.4 (UL128 and UL130 in HCMV) from the unrelated RhCMV strain 180.92, BAC-cloned 68-1 was partially repaired resulting in clone RhCMV 68-1.2 which exhibits broader cell tropism (25). However, the 68-1.2 RhCMV genome sequence still differed significantly from the sequence of low passage RhCMV isolates due to additional mutations that were likely acquired during the prolonged tissue culture of the original 68-1 isolate. The inverted segment in the U_L_-homologous region of 68-1 RhCMV was recently re-examined by amplifying and sequencing DNA from the original urine sample used for virus isolation in 1968 (38). This work revealed the sequence of genes deleted in the U_L_ region upon later passage of 68-1 including the homologs of UL128 and UL130 which showed substantial sequence variation compared to the corresponding genes of 180.92 used in the repaired 68-1.2. This is likely due to significant polymorphism across strains for these genes in RhCMV (39). To create a BAC that most closely resembles a clone of the original 68-1 primary urine isolate we therefore synthesized the entire gene region containing the inverted and missing genes in the U**_L_** region in two overlapping fragments that were then inserted into RhCMV 68-1.2 by homologous recombination (**Fig. 1**). Subsequently, we used *en passant* recombination to repair all point mutation resulting in truncated ORFs as well as a nonsynonymous point mutation in Rh164 (UL141). Finally, we deleted a transposon from Rh167 (O14) that was inadvertently acquired during the construction of the RhCMV 68-1.2 BAC. We confirmed the correct sequence of our BAC by restriction digest and next generation sequencing (NGS) and termed the final construct full length RhCMV (FL-RhCMV).

**Figure 1:**
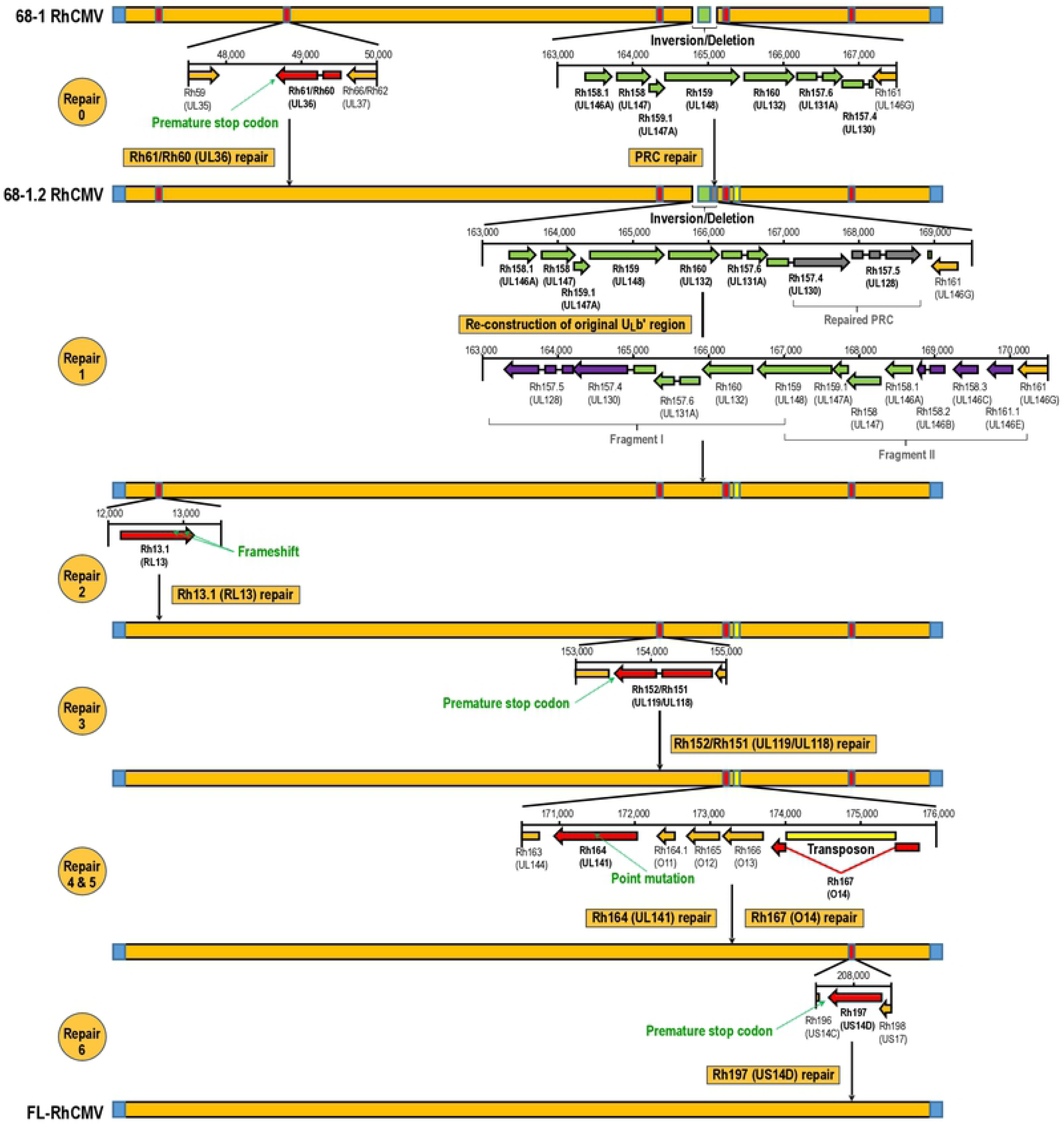
Construction of FL-RhCMV. The schematic depicts the repair steps performed to generate FL-RhCMV. Unaltered ORFs and the unmodified viral genome are shown in orange while the terminal repeats are indicated in blue. ORFs containing known mutations that were repaired in this study are highlighted in red. Genes contained in the acquired inversion in the ULb’ region are shown in green, while genes lost in 68-1 but re-inserted into the genome during the repair are highlighted in purple. The transposon picked up during the generation of 68-1.2 RhCMV is highlighted in yellow and the 180.92 RhCMV PRC members used in the construction of 68-1.2 RhCMV are marked in grey. Repair 0: The frameshift resulting in a premature stop codon in Rh61/60 of 68-1 RhCMV was repaired and the two missing PRC members (Rh157.4 and Rh157.5) were inserted to generate 68-1.2 RhCMV as described previously (25). <u>Repair 1</u>: Two DNA fragments combined spanning 6.9kb corresponding to the genomic sequence of the ULb’ homologous region in the circulating virus originally isolated from sample 68-1 (38) were synthesized. Three undefined bases in the published nucleotide sequence (KF011492) were taken from the consensus sequence of all sequenced low-passage RhCMV isolates. A synthetic DNA fragment spanning the region upstream of Rh157.5 (UL128) to Rh161 (UL146G) in its original orientation was used to replace the corresponding gene region in 68-1.2 RhCMV. The resulting construct maintains the repaired Rh61/60 gene while also containing the original isolate 68-1 genes Rh157.4 (UL128) and Rh157.5 (UL130) as well as the genes coding for the UL146 homologs Rh158.2, Rh158.3 and Rh161.1. <u>Repair 2</u>: Two previously described frameshift mutations in Rh13.1 (10) were repaired resulting in an intact Rh13.1 ORF. <u>Repair 3</u>: A premature stop codon in the viral Fcγ-Receptor homolog Rh152/151 (10) was repaired restoring the ORF to its original length. <u>Repair 4</u>: A nonsynonymous point mutations in Rh164 (UL141) initially predicted by us was confirmed by sequencing the original urine isolate. Hence, we restored the natural DNA sequence. <u>Repair 5</u>: Full genome sequencing of the RhCMV 68-1.2 BAC revealed that an *E. coli* derived transposon had inserted itself into the Rh167 ORF. The transposon was removed by *en passant* mutagenesis and the intact Rh167 ORF was restored. <u>Repair 6</u>: The US14 homologue Rh197 contained a premature stop codon which was repaired.

To characterize the phylogenetic relationship of the 68-1 derived FL-RhCMV BAC clone to related old world NHP CMV species, we cultured and sequenced new isolates from RhCMV, cynomolgus CMV (CyCMV), Japanese macaque CMV (JaCMV) and baboon CMV (BaCMV) from two different US primate centers. We also performed next generation sequencing on viral DNA isolated from stocks of the extensively characterized RhCMV isolates UCD52 and UCD59 grown on epithelial cells and included these genome sequences into our analysis. For comparison, we included all NHP CMV sequences of complete or mostly complete genomes deposited in GenBank (**Fig. 2**). As expected, FL-RhCMV clustered with all other RhCMV isolates and was more distantly related to the CMVs from other NHPs, with the evolutionary relationship of CMV species tracing the evolutionary relationship between their corresponding host species.

**Figure 2:**
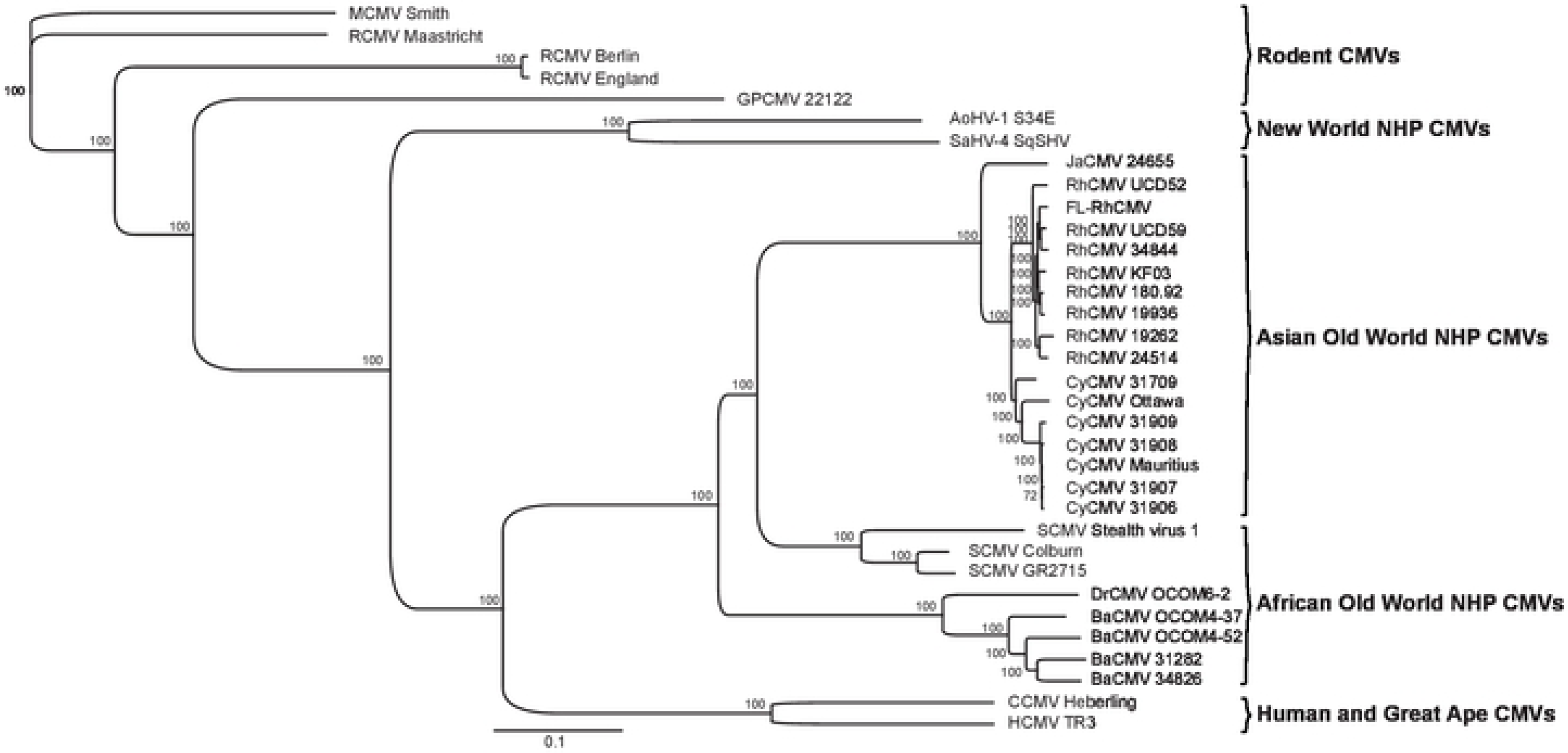
Sequence relationship of FL-RhCMV with NHP CMVs. A phylogenetic tree for FL-RhCMV and rodent and primate CMVs was constructed based on full genome alignments using the Geneious Prime Tree Builder application. Sequences previously published include RhCMV 180.92 (35) as well as the RhCMV isolates 19262, 19936 and 24514 and the Cynomolgus CMV isolates 31906, 31907, 31908 and 31909 (39). We also included the published sequences for the CyCMV strains Ottawa (41) and Mauritius (42), the simian (African green monkey) CMV isolates Colburn (82), GR2715 (40) and stealth virus 1 (83) as well as the BaCMV strains OCOM4-37 (84) and OCOM4-52 (85) and the DrCMV strain OCOM6-2 (85). For comparison we included the HCMV TR3 strain (34), the chimpanzee CMV strain Heberling (86) and the only two complete genome sequences of new world NHP CMVs, Aotine betaherpesvirus 1 (AoHV-1) strain S34E (87) and Saimiriine betaherpesvirus 4 (SaHV-4) strain SqSHV (88). New genome sequences included in this alignment are as follows: the two RhCMV isolates 34844 and KF03, the CyCMV isolate 31709, the Japanese macaque CMV JaCMV 24655 and the two baboon CMVs BaCMV 31282 and 34826. These CMVs were isolated from fibroblast co-cultures of urine samples obtained from NHP housed either at the Oregon National Primate Research Center (ONPRC) or the Tulane National Primate Research Center (TNPRC). Also included in the alignment are the genomic sequences of the previously published RhCMV isolates UCD52 and UCD59 that originated at the UC Davis National Primate Research Center (29, 30). The rodent CMVs include the rat CMV (RCMV) isolates Maastricht (89), England (90) and Berlin (91), the guinea pig CMV (GPCMV) isolate 22122 (92) and the murine CMV (MCMV) strain Smith (93), which was used as an outgroup.

To ensure that FL-RhCMV contained the full ORFeome of all presently confirmed and predicted viral genes of circulating RhCMV strains, we compared the full annotation of FL-RhCMV with that of other old world NHP CMVs (35, 40–42). All RhCMV genomes lack an internal repeat sequence so that the genomic regions corresponding to the unique long (U_L_) or the unique short (U_S_) coding regions are fixed in a given orientation whereas HCMV and ChCMV genomes can freely switch between four isomeric forms (**Supplementary Fig. 1**). Interestingly, the U_S_-homologous region of BaCMV and Drill CMV (DrCMV) is fixed in the opposite orientation compared to RhCMV consistent with an isomer fixation event independent from the RhCMV lineage indicating that single isomers were fixed during the evolution of old world NHP CMVs on more than one occasion. A closer analysis of the genomes revealed that all primary RhCMV isolates without obvious deletions or inversions are predicted to contain the exact same ORFs in the same order. No strain-specific ORFs were identified based on our previously established RhCMV annotation (10). FL-RhCMV contains all ORFs found in other RhCMVs indicating that the full genome content has been restored (**Fig. 3**). A closer examination of full genome alignments of all known old world NHP CMV genome sequences additionally allowed us to further refine our previously established annotations with changes largely comprising reannotations of start codons and splice donor- and acceptor sites (**Supplementary Table 1**). Comparing the viral ORFeomes across old world NHP CMV species revealed a very high degree of conservation in the entire lineage of viruses so that the entire RhCMV annotation can almost seamlessly be transferred to all related species. While our results are based on comparative genomics and hence need to be confirmed experimentally by mass spectrometry or ribozyme profiling, it is interesting to note that most ORFs that differ between NHP CMV species are due to gene duplication events that occurred in six different loci across the genome (**Supplementary Fig. 2-7**). Taken together we conclude that the FL-RhCMV clone we engineered is likely a representative of the genomes contained in the original 68-1 isolate.

**Figure 3:**
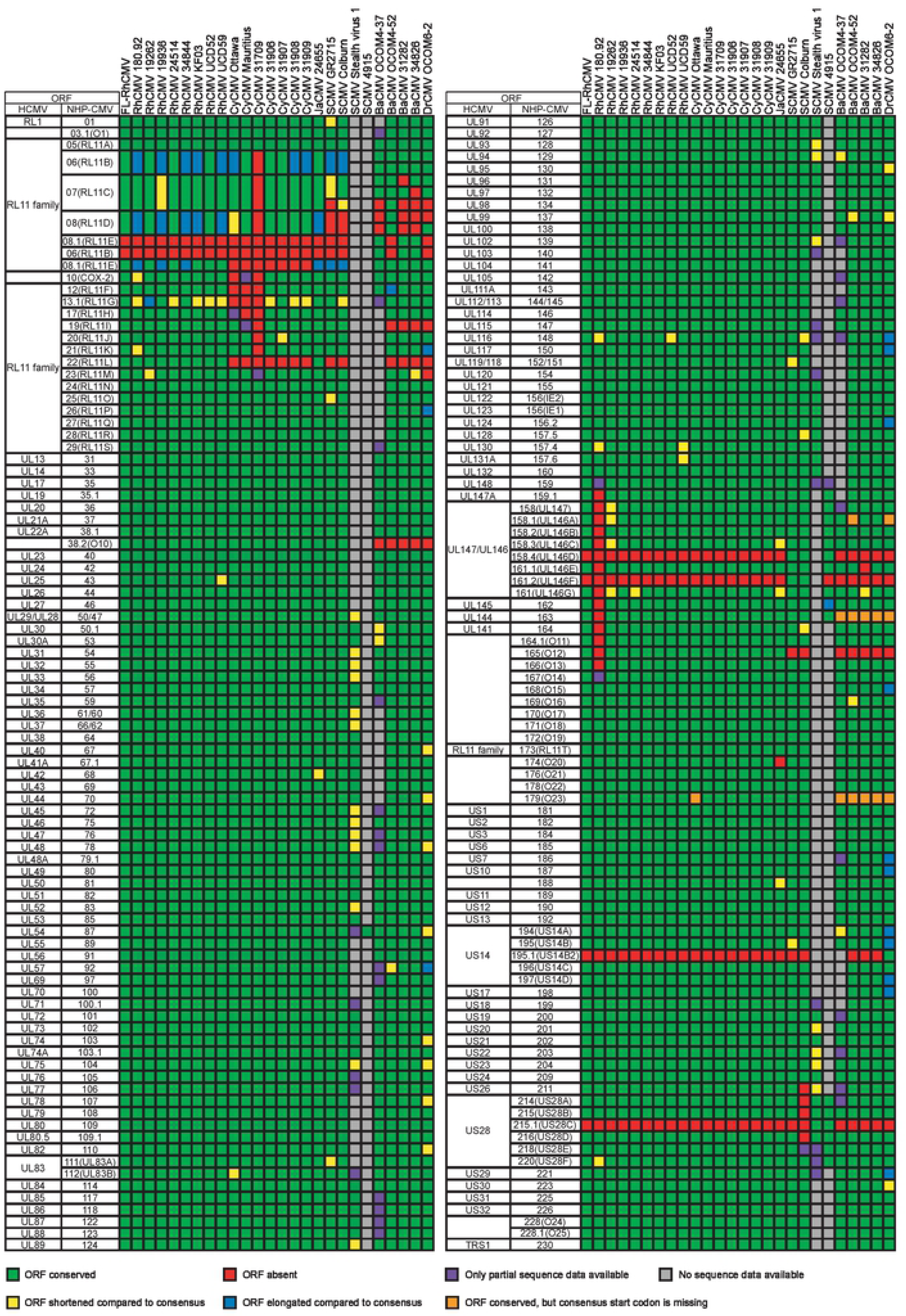
Viral ORFs contained in FL-RhCMV compared to other NHP CMVs. Full genome annotations of all listed old world NHP CMVs are shown. The leftmost column indicates the HCMV nomenclature for CMV encoded genes. Each ORFs that has a defined orthologue in HCMV and old world NHP CMVs is marked. If an orthologue cannot be clearly identified, the homologous gene family is given. The second column identifies the old world NHP CMV nomenclature. The same ORF nomenclature is used across all shown species, with the first or the first two letters corresponding to the host species (e.g. Rh for rhesus macaque). The virus strain analyzed is indicated. Green boxes indicate ORFs present in a particular strain, whereas red boxes indicate ORFs that are absent. Frameshifts or point mutations leading to shortened or elongated ORFs are highlighted in yellow or blue, respectively. Grey boxes indicate absent ORFs due to missing genome sequence information whereas ORFs with partial sequences are highlighted in purple. Orthologous ORFs that lack a conserved canonical start codon in some strains are highlighted in orange.

### *In vitro* characterization of FL-RhCMV

As we have reported earlier (10), one of the ORFs frequently mutated in passaged RhCMV and other old world NHP CMV isolates is the RL11 family member Rh13.1 (**Fig. 3**). This ORF is homologous to HCMV RL13 which is often lost or mutated upon tissue culture passage of HCMV (32). Loss of a functional RL13 protein was shown to result in more rapid cell to cell spread in tissue culture suggesting that RL13-deficient HCMV mutants have a substantial growth advantage *in vitro* (43). Loss of RL13 could be prevented in HCMV by conditional expression of RL13 mRNA under the control of a tet operator (tetO) and growth in tet-repressor (tetR)-expressing fibroblasts (43). As we have repaired this ORF and likely restored its function during the construction of FL-RhCMV we wanted to examine whether conditional expression of Rh13.1 would similarly affect the spread of FL-RhCMV in tissue culture. Hence, we inserted tandem tetO sequences 131 bp upstream of the Rh13.1 start codon and transfected the resulting FL-RhCMV/Rh13.1/tetO BAC DNA into telomerized rhesus fibroblasts (TRFs) expressing tetR (44). The cells were overlaid to prevent cell-free spread and upon recovery of virus we measured viral plaques sizes after 18 days. As a control, we included a FL-RhCMV in which the Rh13.1 ORF had been deleted (FL-RhCMVΔRh13.1). The development of plaques was severely impeded in TRFs transfected with FL-RhCMV or FL-RhCMV/Rh13.1/tetO (**Fig. 4A, B**). In contrast, FL-RhCMVΔRh13.1 spread rapidly in TRF and expression of the tetR led to a partial rescue of plaque formation by FL-RhCMV/Rh13.1/tetO (**Fig. 4A, B**).

**Figure 4:**
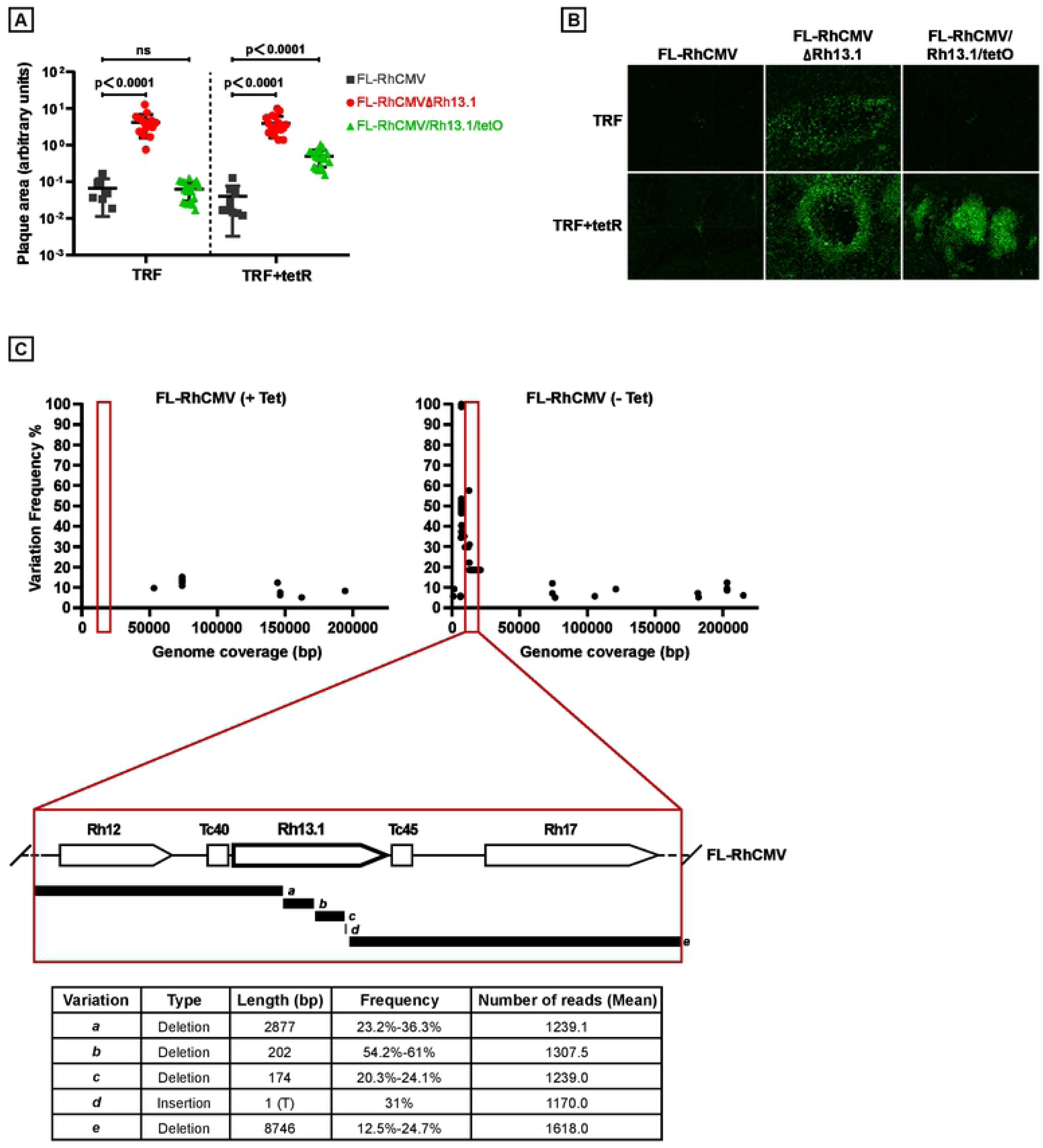
Conditional expression of the RL13 homolog Rh13.1 results in reduced spreading and genomic rearrangements. A) Deletion or reduced expression of Rh13.1 results in increased plaque size. Telomerized rhesus fibroblasts (TRF) or TRF expressing the tet-repressor (tetR) were transfected with the indicated BACs. All recombinant BACs were engineered to express GFP from a P2A linker after UL36 (6). 18 days later plaque sizes were visualized by GFP expression, and measured using ImageJ. Statistical significance was determined using an ordinary one-way ANOVA test with a p-value significance of <0.05. B) Representative images of the GFP positive plaques produced by the indicated constructs on either TRFs or TRFs expressing the tetR are shown. C) Genetic instability of the genomic region surrounding the Rh13.1. Top: The position and relative frequencies of single nucleotide changes were determined by NGS within the genomes of FL-RhCMV/Rh13.1apt after two passages *in vitro* in the presence or absence of tetracycline. The lower panel depicts the positions of deletions/insertions of multiple nucleotides. Frequencies for each deletion are given as percentages of all reads analyzed. Since the short reads generated by the Illumina platform do not cover the entire Rh13.1 locus it is not possible to determine which deletions co-occurred in individual viral genomes resulting in combined frequencies of >100%.

As an alternative approach to conditionally express Rh13.1 we explored the use of aptazyme riboswitches mediating the tetracycline dependent degradation of mRNAs *in cis* (45). We inserted the Tc40 aptazyme sequence upstream and the Tc45 aptazyme sequence downstream of the Rh13.1 coding region in FL-RhCMV and monitored the stability of Rh13.1 and the surrounding genomic region by NGS upon recovery and propagation of virus in the presence or absence of tetracycline. FL-RhCMV/Rh13.1/apt grown in the absence of tetracycline displayed multiple mutations and deletions in this genomic region as early as passage 2 (**Fig. 4C**). In contrast, by activating the aptazyme using tetracycline we were able to generate virus stocks that contained an intact Rh13.1 sequence (**Fig. 4C**). These data are consistent with Rh13.1 being selected against in FL-RhCMV similar to selection against RL13 in HCMV because these homologous proteins impede spread in tissue culture. We further conclude that mutations in the Rh13.1 homologs found in many old world NHP CMV genomes (**Fig. 3**) are due to rapid tissue culture adaptations whereas the parental isolates likely contained an intact ORF. Thus, Rh13.1 and its homologs are preserved *in vivo*, but are selected against *in vitro*.

It was previously shown that repair of the PRC increased the ability of RhCMV 68-1.2 to infect epithelial and endothelial cells without affecting growth characteristics in fibroblasts (25). Similarly, growth characteristics of FL-RhCMV/Rh13.1/apt were comparable to that of 68-1 and PRC-repaired 68-1.2 in rhesus fibroblasts with respect to kinetics and peak titers in a multistep growth curve (**Fig. 5A**). Since these comparable growth kinetics were observed in the absence of tetracycline induced Rh13.1 mRNA degradation, we conclude that while the presence of Rh13.1 does effect virus cell to cell spread after transfection of BAC DNA (**Fig. 4A, B**), we cannot observed a phenotype in the context of a multistep-growth curve when starting with infectious virus and without overlaying the cell monolayer to prohibit cell free spread. Since the PRC is important for entry into non-fibroblast cells (25) we quantified infection levels of 68-1, 68-1.2 and FL-RhCMV upon entry into rhesus retinal epithelial (RPE) cells or primary rhesus fibroblasts using flow cytometry. When normalized to infected fibroblasts, 68-1 showed a strongly reduced ability to enter RPE compared to 68-1.2 (**Fig. 5B**) consistent with previous reports (25). In contrast, a FL-RhCMV vector carrying an SIVgag insert replacing Rh13.1 (FL-RhCMVΔRh13.1gag) displayed an increased ability to enter RPE cells compared to 68-1. However, infection rates on RPE cells with FL-RhCMV were consistently lower compared to RhCMV 68-1.2 in multiple independent experiments. In HCMV, strain specific differences in tropism can arise from alterations in the levels of both the PRC and the gH/gL/gO trimeric receptor complex which can be caused by genetic sequence variations or altered mRNA expression levels of the proteins in each complex (46–48). Intriguingly, increased infection rates on epithelial cells have been reported for the PRC-repaired HCMV strain AD169 compared to PRC-intact low-passage isolates (49), a result very reminiscent of our data. This difference was determined to be due to the absence of the UL148 glycoprotein in AD169, a protein that will reduce PRC levels in favor of the trimeric gH/gL/gO complex on the virus membrane (50). Similarly, mRNA expression levels of the UL148 homologue Rh159 late during infection were higher in FL-RhCMV compared to both 68-1 and 68-1.2 (**Fig. 5C**). Since this gene is located within the genomic region that was inverted in 68-1, it is likely that it was put out of context of its original regulatory DNA elements, resulting in altered mRNA expression levels. Consistent with this explanation, examination of the mRNA expression levels of the late Rh137 (UL99, pp28) gene not encoded within the acquired inversion did not show any significant differences across the examined strains (**Fig. 5C**). We previously demonstrated that Rh159 is an ER-resident glycoprotein that intracellularly retains NK cell activating ligands, a function that is not shared with UL148 (51). However, these observations do not rule out a role of Rh159 for PRC expression and cell tropism. While further work will be required to establish this role, our results indicate that FL-RhCMV is remarkably similar to low passage clinical isolates of HCMV with respect to growth in tissue culture, tissue tropism and genetic stability *in vitro*.

**Figure 5:**
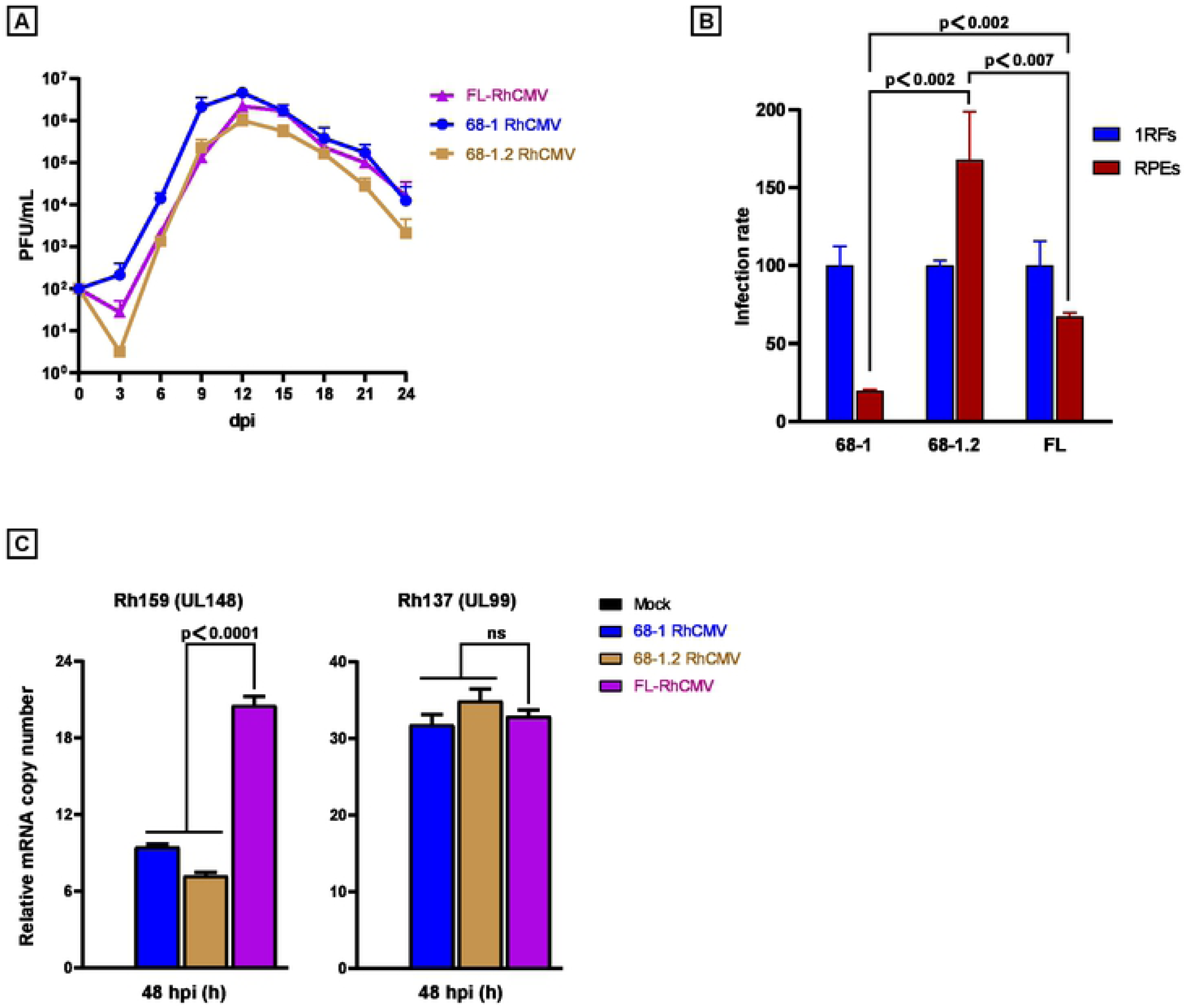
Growth of FL-RhCMV *in vitro*. A) Multistep growth curve of FL-RhCMV in fibroblasts indicates growth comparable to 68-1 and 68-1.2 RhCMV. Primary rhesus fibroblasts were infected with RhCMV 68-1, 68-1.2 and FL-RhCMV/Rh13.1/apt at an MOI of 0.01 on day 0. Cell culture supernatants were harvested on the indicated days and virus titers were determined by TCID50. Results of two biological repeats titrated in duplicate are shown and the standard error of the mean (SEM) is indicated by error bars. B) FL-RhCMV shows increased infection of epithelial cells compared to 68-1 RhCMV. Primary rhesus fibroblasts or rhesus retinal epithelial cells (RPE) were infected with MOIs of 0.3 or 10, respectively, and all experiments were performed in triplicates. After 48 hours post infection, cells were harvested, fixed, permeabilized, stained with a RhCMV specific antibody (52) and analyzed by flow cytometry. Statistical significance was shown using an unpaired t-test with a p-value significance threshold of <0.05. C) Rh159, the RhCMV homologue of UL148, is upregulated in FL-RhCMV. Relative mRNA copy numbers of Rh159 (UL148) and Rh137 (UL99) were determined by quantitative reverse transcription polymerase chain reaction (qRT-PCR) using specific probes. The data shown represent the mean of triplicate repeats (+/- SEM). Unpaired student t-tests with a p-value significance threshold of <0.05 were performed to show statistical significance in both graphs comparing FL-RhCMV/Rh13.1/apt to either 68-1 RhCMV or 68-1.2 RhCMV at 48 hpi.

### Kinetics and magnitude of infection by FL-RhCMV is similar to wildtype RhCMV

It was reported previously that different from the low passage isolates UCD52 and UCD59, 68-1 RhCMV displayed severely reduced viral genome copy numbers in plasma, saliva and urine in RhCMV-seronegative RM after experimental subcutaneous (s.q.) infection (26). Concordantly, we observed significant plasma viremia in three female RhCMV-naïve pregnant RM infected intravenously (i.v.) with 1×10^6^ pfu of RhCMV UCD52, 1×10^6^ pfu of RhCMV UCD59, and 2×10^6^ TCID_50_ of RhCMV strain 180.92 (**Fig. 6A**) consistent with previous reports (11). Viral genome copy numbers of approximately 10^5^ copies/ml blood were detected by qPCR in all three animals between days 7 to 21, declining thereafter. Similar kinetics of infection and peak viremia were measured in three male RhCMV-naïve RM infected i.v. with Rh13.1-intact FL-RhCMV/Rh13.1/apt grown in the presence of tetracycline to maintain genome integrity during virus stock production (**Fig. 6B**). Since both experiments showed virtually the same development and progression of plasma viremia after i.v. inoculation (**Fig. 6C**), we conclude that *in vivo* replication of FL-RhCMV is comparable to that of low passage RhCMV isolates.

**Figure 6:**
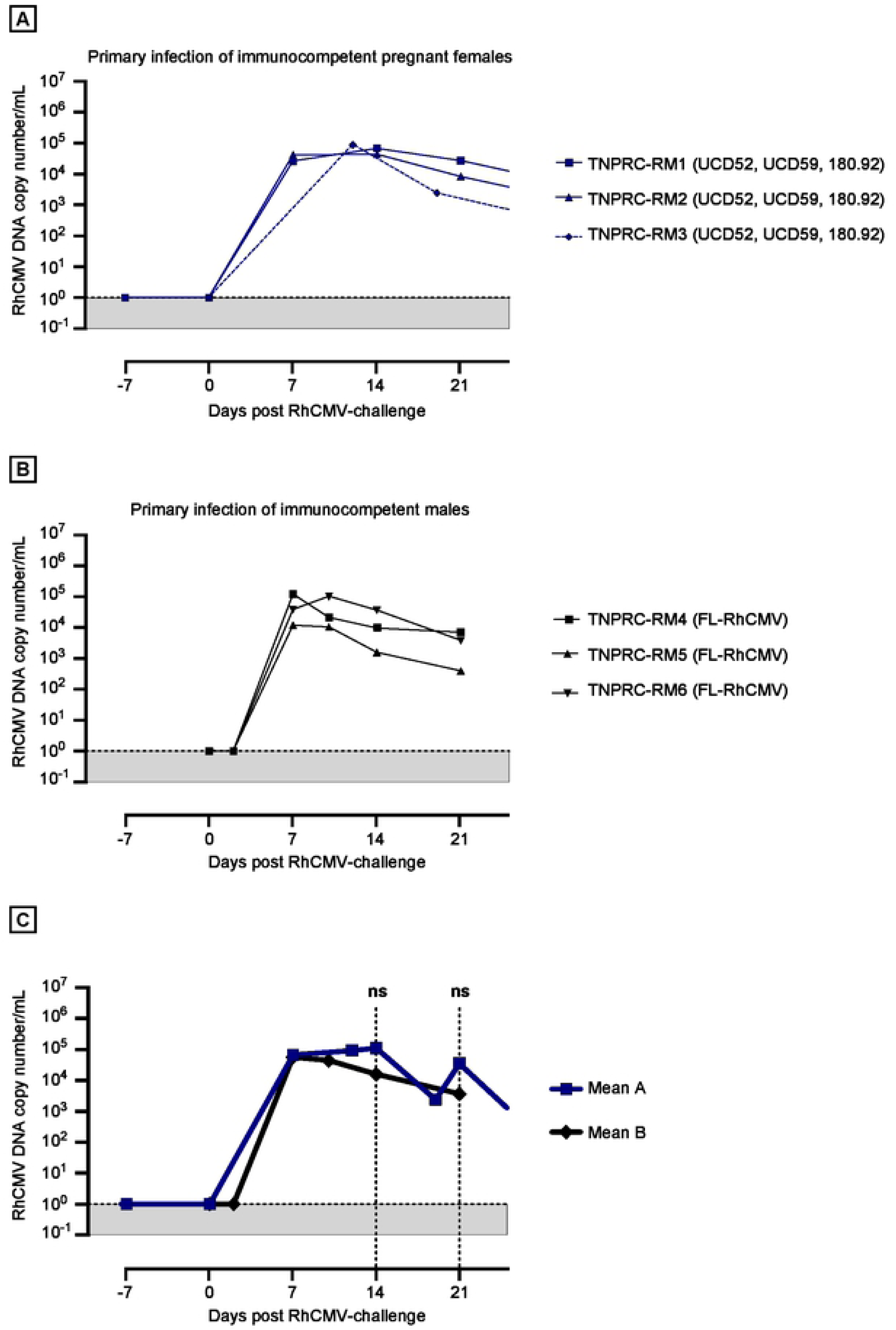
*In vivo* replication of FL-RhCMV is similar to low passage isolates in RhCMV-negative animals. A) Replication of RhCMV isolates UCD52, UCD59 and 180.92 in RhCMV seronegative RM. Plasma RhCMV DNA load in three RhCMV-seronegative pregnant female RM inoculated i.v. with 2 x 10^6^ TCID50 RhCMV 180.92, 1×10^6^ PFU RhCMV UCD52, and 1×10^6^ PFU RhCMV UCD59. The plasma viral load (PVL) in each RM was determined at days 7, 14 and 21 using qPCR for the exon 1 region of the immediate early gene as described previously (11, 72). B) Replication of FL-RhCMV in RhCMV-seronegative RM. 1.79×10^6^ PFU of FL-RhCMV/Rh13.1apt were inoculated i.v. into three RhCMV-seronegative male RM and the PVL was determined at the indicated days by qPCR of the RhCMV gB gene as described previously (71). The PVL for each RM is shown. C) The mean PVL for all animals infected in A) and B) are shown in comparison and compared at days 14 and 21 by non-parametric Mann-Whitney test. The data indicates that FL-RhCMV can induce a PVL comparable to commonly used virulent RhCMV isolates.

### *In vivo* dissemination of FL-RhCMV-derived viral vectors

A central goal of our research is to use the RhCMV animal model for the development of CMV-based vaccine vectors (12). We recently reported that deletion of the pp71-encoding RhCMV gene Rh110 resulted in reduced dissemination and lack of shedding of 68-1-derived vaccine vectors (20). Nevertheless, SIV-antigen expressing vaccines based on these live-attenuated vector backbones maintained the ability to control highly virulent SIV_mac239_ upon challenge (19). However, since 68-1 lost its homologs of the PRC subunits UL128 and UL130 as well as homologs of the viral UL146 family of CXC chemokines it was conceivable that these deletions contributed to viral attenuation. To determine the impact of these gene deletions on viral dissemination we generated viral vectors based on either FL-RhCMV or FL-RhCMV lacking the homologs of UL128, UL130 and all members of the UL146 family. As PCR- and immunological marker we selected a fusion protein of six *M. tuberculosis* antigens that was recently used to demonstrate protection against *Mtb* challenge with 68-1-based vaccine vectors (14). As antigen-insertion site we replaced the Rh13.1 gene, thus increasing vector stability while using the Rh13.1 promoter to drive antigen expression.

As we have shown previously (52), *in vivo* dissemination of 68-1 RhCMV is observed in RhCMV-seronegative animals. Hence we assigned six CMV-naïve RM and inoculated three with FL-RhCMVΔRh13.1/TB6Ag and three with FL-RhCMVΔRh13.1/TB6AgΔRh157.4-5ΔRh158-161. Subsequently, we took the animals to necropsy at 14 dpi to systematically measure viral genome copy numbers in tissue samples using nested PCR as described previously (20, 52). While both recombinants resulted in significant viral accumulation at the injection sites and the nearest draining lymph node, FL-RhCMV genomic DNA was highly abundant in many of the tissues examined (**Fig. 7A**). In contrast, FL-RhCMV lacking the UL128, UL130 and UL146 homologs displayed significantly reduced spreading beyond the initial site of replication (**Fig. 7A**). Solely tissue samples from the spleen retained notable viral copy numbers for the deletion mutant, although at significantly reduced levels compared to FL-RhCMV (**Fig. 7B, C**), whereas dissemination to and replication in most other tissues was almost completely abrogated. The results obtained with FL-RhCMVΔRh13.1/TB6AgΔRh157.4-5ΔRh158-161 and FL-RhCMVΔRh13.1/TB6Ag are consistent with previous observations for 68-1 RhCMV (20, 52) and UCD52 and UCD59 (26), respectively. These data also suggest that RhCMV vectors with 68-1 like configuration are attenuated *in vivo*.

**Figure 7:**
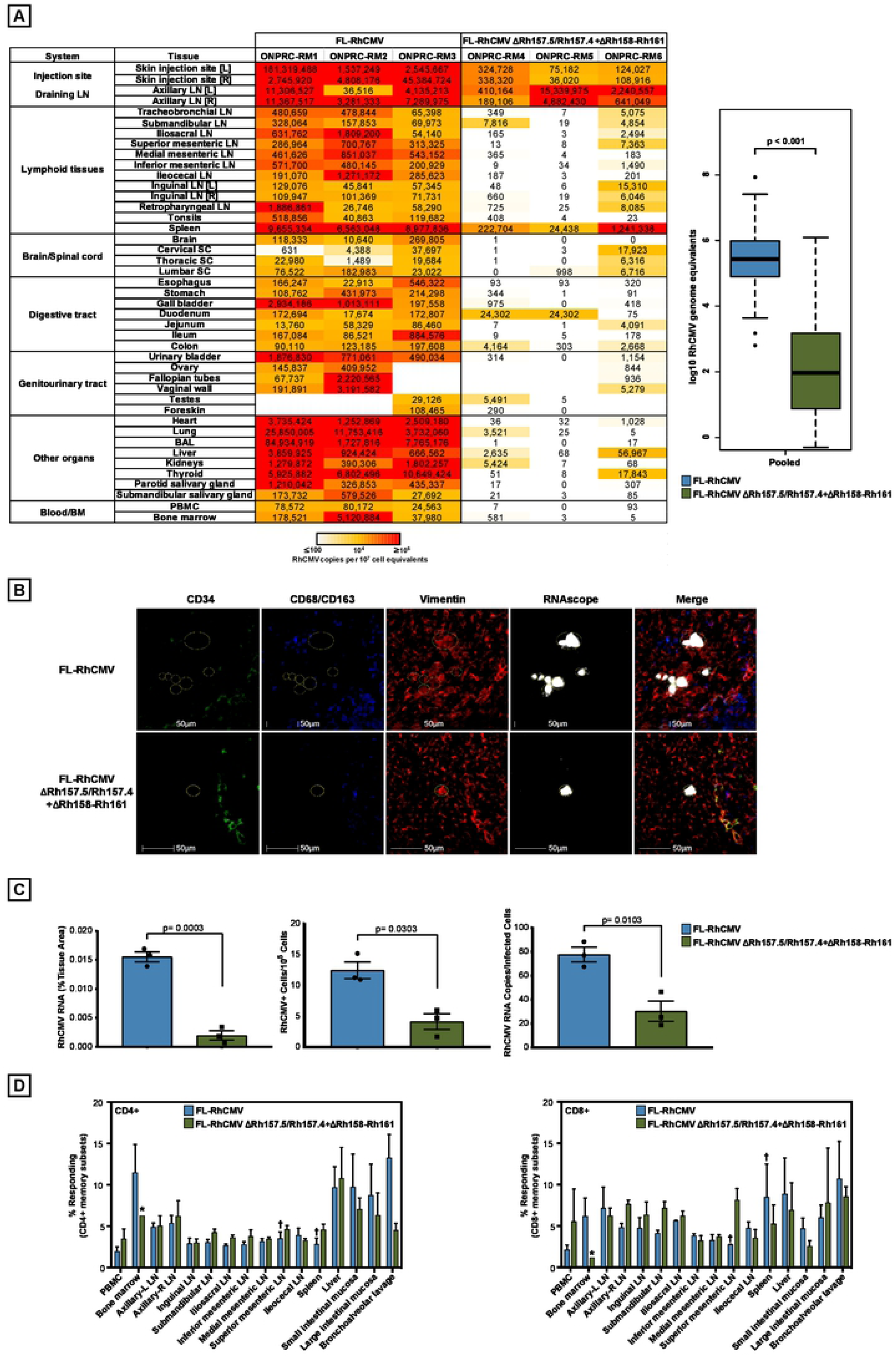
Spreading of FL-RhCMV *in vivo*. A) Tissue genome copy numbers of FL-RhCMV. Three RhCMV-naïve RM (RM1-RM3) were inoculated with 10^7^ PFU FL-RhCMVΔRh13.1/TB6Ag while another three RhCMV-naïve RM (RM4-RM6) were inoculated with 10^7^ PFU of FL-RhCMVΔRh13.1/TB6AgΔRh157.4-5ΔRh158-161. All 6 RM were necropsied at day 14 post-infection and viral genome copy numbers per 10^7^ cell equivalents were determined in the indicated tissues using ultra-sensitive nested qPCR specific for TB6Ag. Statistical analysis was performed using a two-sided Wilcoxon tests (unadjusted p values < 0.05) excluding all tissues at the injection site and the nearest draining lymph nodes to detect significant differences in dissemination. (B) *In situ* immunofluorescence phenotyping of cells expressing RhCMV RNA was performed by multiplexing RNAscope *in situ* hybridization with antibody detection of cellular markers specific for myeloid/macrophage cells (CD68/CD163), endothelial cells (CD34), and mesenchymal cells (vimentin) in the spleen of macaques inoculated with either FL-RhCMVΔRh13.1TB6Ag (FL-RhCMV) or FL RhCMVΔRh13.1/TB6AgΔRh157.4-5ΔRh158-161. The majority of cells inoculated with the FL-RhCMV were vimentin+ CD34- CD68/CD163-, indicating they were of mesenchymal origin. (C) To quantify differences in RhCMV infection and expression levels in macaques inoculated with either FL-RhCMV or FL RhCMVΔRh13.1/TB6AgΔRh157.4-5ΔRh158-161, we used three independent quantitative approaches in the HALO image analysis platform from Indica Labs: i) the percent area of the tissue occupied by infected cells, ii) the number of infected cells per 10^5^ cells, and iii) an estimate of RhCMV viral RNA copy number per infected cell. Statistical significance was calculated using an unpaired t-test. D) Tissue distribution of Tb6Ag insert–specific CD4^+^ and CD8^+^ T cell responses elicited by FL-RhCMVΔRh13.1TB6Ag versus FL RhCMVΔRh13.1/TB6AgΔRh157.4-5ΔRh158-161 vectors. Flow cytometric ICS (CD69, TNF-α and/or IFN-γ readout) was used to determine the magnitude of the CD4^+^ and CD8^+^ T cell responses to peptide mixes corresponding to the six Mtb antigens contained in the TB6Ag-fusion (Ag85A, ESAT-6, Rpf A, Rpf D, Rv2626, Rv3407). Mononuclear cells were isolated from the indicated tissues from three RhCMV-naïve RMs inoculated with 10^7^ PFU FL-RhCMVΔRh13.1TB6Ag (blue bars) and three RMs inoculated with 10^7^ PFU FL-RhCMVΔRh13.1/TB6AgΔRh157.4-5ΔRh158-161 (green bars) and all RM taken to necropsy at either 14 or 15 days post infection. Response comparisons per tissue are shown as the mean + SEM percentage of T cells specifically responding to the total of all peptide mixes (background subtracted) within the memory CD4^+^ or CD8^+^ T cell compartment for each tissue (n=3 per tissue, unless otherwise noted by * n=1 or † n=2).

To determine the cell types infected *in vivo* by FL-RhCMV during primary lytic replication, we performed RNAscope *in situ* hybridization (ISH) in combination with immunohistochemistry for cellular markers on spleen tissue obtained from the same animals. Consistent with the high genome copy numbers observed, FL-RhCMVΔRh13.1TB6Ag was rapidly detected in tissue sections of the spleen, the large clusters of infected cells primarily localized within the white pulp with fewer individual infected cells within the red pulp (**Fig. 7B, C**). In contrast, the deletion mutant could only be detected very sparsely in a few infected cells across the examined tissue, localized primarily within the white pulp with sporadic rare viral RNA+ cells found within the red pulp. Co-staining with cellular markers identified the infected cells as vimentin-positive mesenchymal cells such as fibroblasts, whereas we did not find RhCMV RNA in CD34^+^ hematopoetic stem cells or CD68/CD163-positive macrophages commonly associated with CMV latent infection. While this does not exclude the possibility that FL-RhCMV can infect other cells types, it indicates that the vast majority of cells infected in the spleen during the initial acute viremia after infection of naïve RM are in the connective tissue. Nevertheless, while the viral loads of the two different vectors differ substantially across naïve RM, they both elicited similar frequencies of TB6Ag-specific CD4^+^ and CD8^+^ T-cell responses in all examined tissues (**Fig. 7D**). This observation is consistent with previously reported findings that, above a given threshold, T cell responses are largely independent of viral replication *in vivo* and with the reported immunogenicity of 68-1-based vectors (14, 15, 20, 53).

### RNAseq analysis of *in vivo* tissue samples identifies multiple viral transcript that are highly expressed across tissues

The high genome copy numbers measured in several tissues of FL-RhCMVΔRh13.1/TB6Ag-inoculated RM at 14dpi provided an opportunity to monitor viral gene expression from a fully characterized viral genome by RNAseq analysis *in vivo*. Total RNA was isolated from the lung, the axillary lymph node (ALN), the parotid salivary gland (PSG) and the submandibular salivary gland (SSG) as these samples showed high viral genome copy numbers (**Fig. 7A**). For comparison, we infected primary rhesus fibroblast at an MOI of 5 with FL-RhCMVΔRh13.1/TB6Ag and harvested total mRNA at 8, 24 and 72 hpi representing immediate early, early and late times post infection. While the average number of reads/sample were comparable between the *in vitro* (average of 86,013,721) and *in vivo* (average of 107,502,852) RNA samples, the ratio of viral/host reads was much higher *in vitro*, particularly at late times of infection, an entirely expected result as a much lower number of cells are infected in our *in vivo* samples (**Supplementary Fig. 8**). The absolute number of reads aligning to the annotated RhCMV ORFs for all *in vitro* and *in vivo* samples can be found in **Supplementary Table 2**. Analysis of *in vitro* samples across the entire FL-RhCMV genome indicated that mRNA expression of all re-introduced genes in the repaired U_L_-region was detectable at all time points, indicating successful restoration of a WT like gene expression cascade (**Supplementary Fig. 9**). Principal component analysis (PCA) on the normalized count matrix of RhCMV transcripts revealed that while the early *in vitro* samples clustered together, the late samples showed an mRNA expression pattern closer to expression profiles obtained from lung and ALN samples (**Fig. 8A**). This is consistent with active viral replication in these tissues at this time point. In contrast, PCA revealed that gene expression profiles of PSG and SSG samples were distinct from the other tissue samples and one another. Importantly, although generated from different outbred animals, viral gene expression patterns from the same tissue source were more closely related across all three RM than across different tissue samples within the same animals. This indicates that viral mRNA expression varies within infected animals depending on the examined tissue.

**Figure 8:**
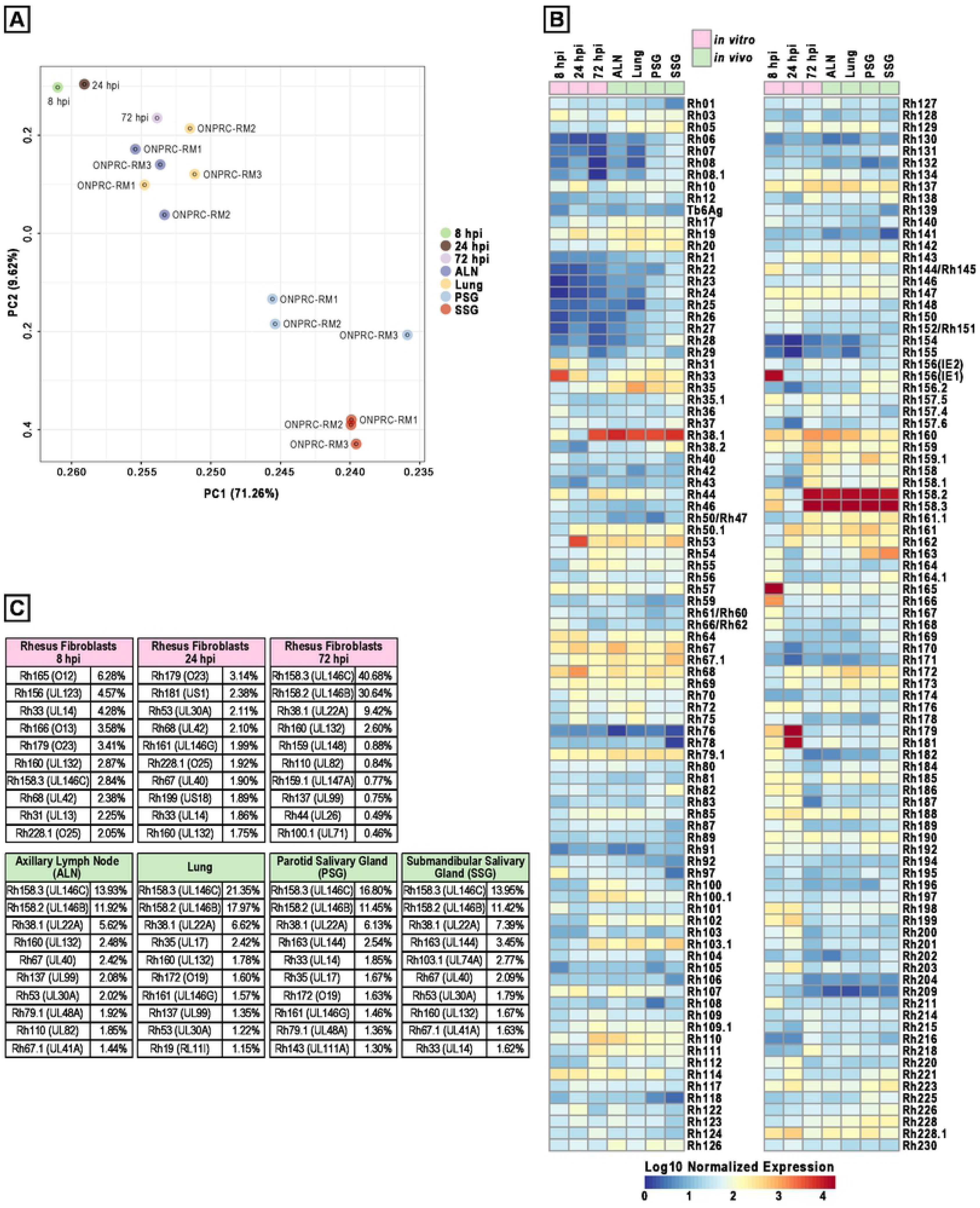
Viral gene expression profile of FL-RhCMV *in vitro* and *in vivo*. A) Comparison of *in vitro* and *in vivo* gene expression profiles by principal component analysis. *In vitro*: Rhesus fibroblasts were infected with an MOI=5 of FLRhCMVΔRh13.1/TB6Ag and the cells were harvested at the indicated times. *In vivo*: RNA-was isolated from indicated tissues of RM1-RM3 described in Fig. 7. Total RNA was isolated from all samples and RNAseq was performed on libraries build from polyA-fractionated RNA using an Illumina HiSeq-2500 next generation sequencer. PCA was done on the combined and quantile normalized expression matrix (see Methods). We observed that PC1 and PC2, shown herein, combined capture over 70% of total variance with distinct sets of co-regulated genes. B) *In vitro* and *in vivo* expression levels of each ORF. Expression levels were normalized between the *in vitro* and *in vivo* samples using quantile normalization (see Methods). C) For all samples analyzed in B) the ten viral ORFs showing the highest mRNA coverage after normalization for ORF size are shown. All values are given as percent of total viral reads mapping to all annotated ORFs normalized for size.

Since expression patterns of the same tissues across animals were comparable, we combined these samples to compare the expression levels of each ORF between tissues and *in vitro* results. Surprisingly, this analysis revealed that some of the most highly expressed ORFs found both at late times post-infection *in vitro* and in all tissues examined *in vivo*, were the soluble chemokine binding protein Rh38.1 (UL22A) as well as two CXC chemokine-like genes of the UL146 family, Rh158.2 (UL146B) and Rh158.3 (UL146C) (**Fig. 8B**). Normalized to the ORF size, of the ten ORFs with the most sequence coverage in each sample (**Fig. 8C**), 80.74% mapped to these three ORFs in rhesus fibroblasts at 72 hpi, 45.94% in Lung, 31.47% in ALN, 34.38% in PSG and 32.76% in SSG. While UL22A is known to be one of the most highly expressed ORFs in HCMV (54), this dominant expression of two UL146 family members had not been observed previously. Interestingly, both UL146-homologs are deleted in 68-1 and were re-inserted during our construction of FL-RhCMV (**Fig. 1**). The abundance of viral transcripts encoding chemokine-binding and chemokine-like proteins suggests that RhCMV interference with the host’s chemokine network is a major immune modulatory strategy during the acute phase of infection.

## Discussion

To generate a RhCMV clone that is representative of a low passage isolate we chose to repair an existing BAC clone instead of cloning a new primary isolate since this would allow us to better compare results obtained with FL-RhCMV to historic data obtained with 68-1 BAC clone-derived recombinants and thus facilitate the identification of viral determinants of tissue tropism, pathogenesis and immune response programming. Using sequence information from the original primary 68-1 isolate (38, 55) and next generation sequencing of multiple primary RhCMV isolates we identified and reverted all mutations that resulted in frameshifts or premature termination codons in predicted ORFs. While we cannot rule out that additional mutations, particularly in non-coding regions, occurred during tissue culture as has been observed in HCMV (46), these cannot be unequivocally distinguished from strain-specific nucleotide polymorphisms and therefore remained unchanged. Since the original isolate likely contained a multitude of molecular clones, our FL-RhCMV is representative of a sub-population present in the 68-1 isolate. Full genome alignments of the old world NHP sequences generated in this study together with sequences previously submitted to GenBank allowed us to refine the genome annotation, enabling more precise genetic engineering of FL-RhCMV derived constructs in the future. Comparative genomics revealed a close conservation of the overall ORFeome across NHP CMV species (**Fig. 3**) while also allowing us to identify differences acquired by individual species during co-evolution with their respective host. These distinct disparities largely consist of gene duplications in only six different loci across the genome and they are reminiscent of a poxvirus adaptation strategy deployed to adapt to antiviral pressure by the immune system known as a genomic accordion (56), albeit on a significantly longer evolutionary timescale. Hence, these gene duplications could be the results of the ongoing arms race between the virus and the host immune defenses. At a minimum, these data enable us to estimate when different CMV gene families entered the NHP CMV lineage and how they adapted over millions of years of co-evolution with their primate host (**Fig. 9**).

**Figure 9:**
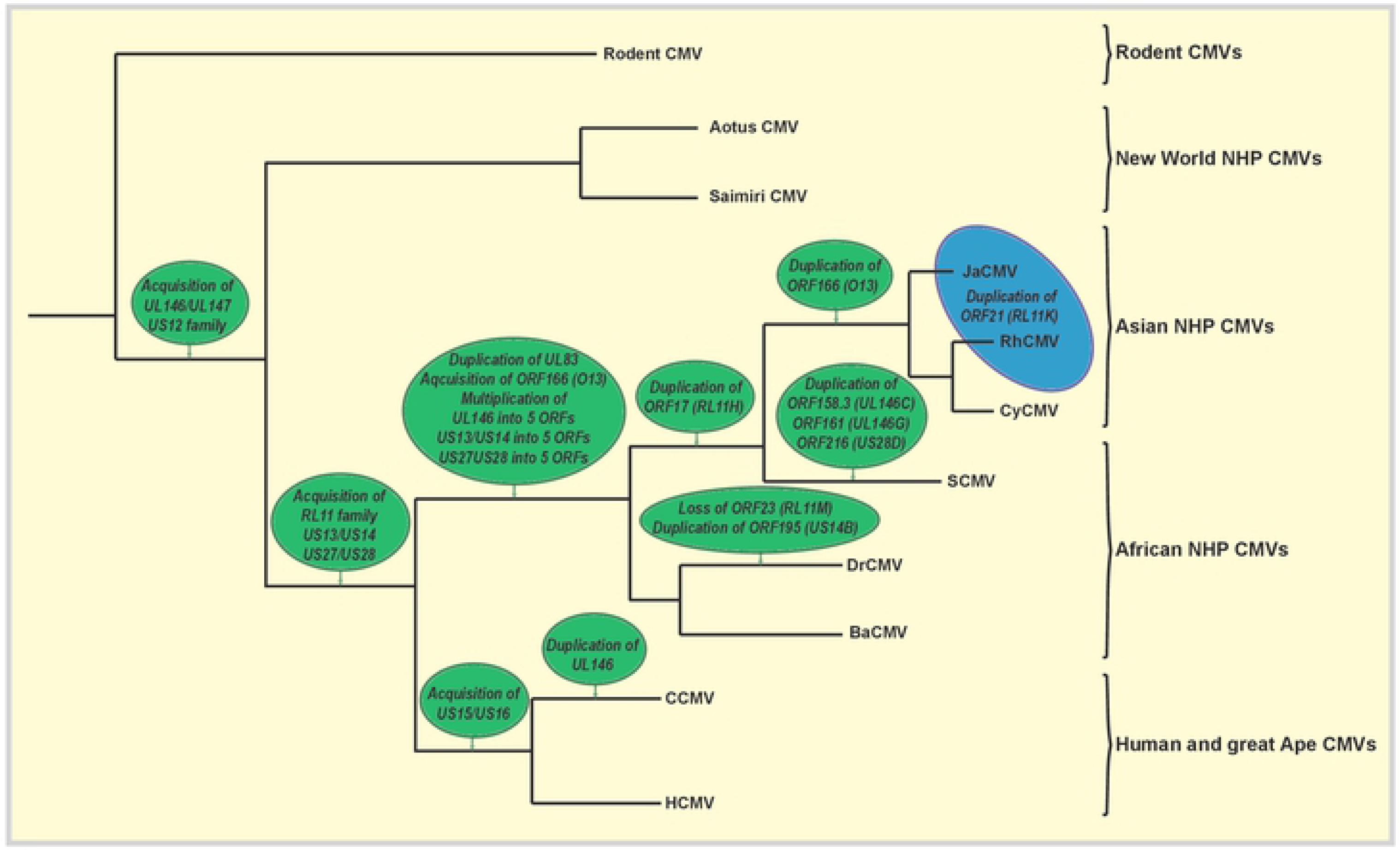
Acquisition of genes and gene families in old world NHP CMVs during co-evolution with their primate hosts. Full genome annotations of old world NHP CMVs across different species allowed for comparative genomics to identify single ORFs or entire gene families that were present in one or several CMV species but absent in others. These differences clustered in six independent loci across the genome. Examination of the phylogenetic relationship of the individual CMV species as well as their host species reveals at which point in time these gene acquisitions and gene duplications occurred. The phylogenetic tree depicted is based on the full genome alignment shown in Fig. 1. The green circles indicate genetic events that took place during the evolution of each species. The blue circle represents the acquisition of RL11K, a gene duplication found in RhCMV and JaCMV but not in CyCMV. Since CyCMV appears to be more closely related to RhCMV than JaCMV by full genome alignment (see Fig.1) this appears counterintuitive, however, phylogenetic alignments of the corresponding macaque host species based on morphology (94), mitochondrial DNA data (95) or Alu elements (96) reveals that Japanese macaques (*M. fuscata*) and rhesus macaques (*M. mulatta*) speciated more recently compared to cynomolgus macaques (*M. fascicularis*).

Another advantage of our chosen repair strategy was that it allowed us to recreate a complete genome at the BAC stage in the absence of the selective forces of *in vitro* tissue culture. Indeed, upon culture of BAC derived Rh13.1-intact FL RCMV in fibroblasts we observed the rapid accumulation of mutations in the gene and the region surrounding the gene. This is strikingly similar to the clinical HCMV isolate Merlin which displayed the same instability in the homologous gene RL13 when an RL13-intact BAC was used for transfection (43). Similar observations have been reported for additional HCMV isolates (32), although a small number of passaged strains appear capable of maintaining an intact RL13 ORF (57). RL13 seems to limit viral spread, particularly in fibroblasts (57) but the exact mechanism of this inhibition is not clear. Rh13.1 belongs to the RL11 family of single transmembrane glycoproteins present in all old world NHP CMVs, as well as great ape and HCMV, but not in CMVs of new world primates (**Fig. 9**). The functional conservation of Rh13.1 and RL13 is surprising since the RL11 family is highly diverse both within a given CMV species and especially when comparing family members between great ape and old world monkey CMVs (10, 40, 58). It thus seems likely that while Rh13.1 and RL13 are selected against *in vitro*, there is a strong selection for their presence *in vivo*. RL13 has been shown to bind to antibody Fc portions (59) and it is thus possible that it serves as an immune evasion protein in the host. Whether this function is conserved in NHP CMVs is currently unknown. To enable the study of Rh13.1-intact vectors, we therefore generated two different tetracycline-regulated systems that allow for the conditional expression of Rh13.1 so that the virus can be grown *in vitro* without selection against Rh13.1 whereas mRNA would be expressed *in vivo* in the absence of tetracycline. Indeed, we observed *in vivo* viremia of Rh13.1-intact FL-RhCMV that was comparable to low passage isolates.

However, we also observed that FL-RhCMV lacking Rh13.1 displayed substantial *in vivo* spread that was significantly more pronounced than a mutant that lacked the UL128 and UL130-homologous subunits of the pentameric complex together with all genes homologous to HCMV UL146 and UL147. These data suggest that Rh13.1-deleted viruses might maintain most of the wildtype characteristics in primary infection of immunocompetent adult animals. We also know that Rh13.1 is not required for the establishment and maintenance of persistent infections since strain 68-1 lacks a functional Rh13.1, yet persists as shown by long-term immune responses and shedding (53). Given a possible function of Rh13.1 in evasion of antibody responses, it would be interesting to compare spreading of Rh13.1 intact and deleted viruses in the presence of anti-CMV antibodies. While inactivation of RL13 generally represents the first tissue culture adaptation step observed in HCMV, this is often followed by the loss of one or multiple members of the PRC (43). Although we did not observed loss of PRC members in FL-RhCMV, this might be due to the limited numbers of passages we examined. Since PRC mutations occurred in 68-1 RhCMV during prolonged tissue culture it is likely that further passaging of FL-RhCMV would result in adaptations akin to HCMV indicating that the overall genome stability and the sequence of adaptation are likely similar across primate CMV species.

The strong attenuation of a “68-1 like” FL-RhCMV lacking homologs of UL128, UL130 and UL146 observed in primary infection suggests that gene deletions in these regions are the likely reason for the previously reported lack of measurable viremia and shedding of 68-1 RhCMV (26). Further studies will be required to determine the individual contribution of pentamer subunits and UL146-related chemokine homologs to viral dissemination and spread. However, it was unexpected that two of the most highly expressed genes, both *in vitro* and *in vivo*, belong to the UL146 family and that these two genes, Rh158.2 (UL146B) and Rh158.3 (UL146C), are deleted in strain 68-1. The UL146 gene family of chemokine like proteins is only found in primate CMVs (**Fig. 9**). Given their homology to chemokines such as interleukin 8 they were likely acquired from the host. HCMV contains two family members: UL146 and UL147. While a single UL147 homologous gene can be found across all primate CMV species with a moderate level of conservation, the UL146-homologs are highly diverse within a CMV species. Moreover, while HCMVs only contain a single UL146 member, the number of genes can vary greatly in other primate CMVs (**Supplementary Fig. 4**). CCMV contains two genes almost equally related to the HCMV UL146 member. However, new world NHP CMVs contain a single UL146 family member that is highly divergent. Conversely, old world NHP CMVs encode five to seven UL146 homologs. Since 68-1 lacks these highly expressed genes, they are not required for the establishment and maintenance of persistent infection but it is possible that these chemokine homologs support viremia and dissemination during primary infection or upon re-infection, a possibility reinforced by the recent observation that inserting the HCMV UL146 protein into MCMV significantly enhances virus dissemination kinetics in infected mice (60).

The *in vitro* and *in vivo* characteristics of FL-RhCMV described here are consistent with this virus being representative of wildtype RhCMV. Based on these observations we anticipate that this recombinant will be useful for RM models of CMV pathogenesis, such as the fetal inoculation model as well as a model of congenital infection (61, 62). In addition, FL-RhCMV can serve as a translational model for the development of live-attenuated vectors derived from clinical isolates of HCMV. As recently reported, our strategy for HCMV-based vectors is to start with a clinical isolate to ensure persistence and then introduce genetic modifications that increase vector safety while maintaining desired immunological features (34). The availability of a complete RhCMV genome will allow us to recapitulate HCMV-vector design strategies and test these designs in RM challenge models for AIDS, tuberculosis and malaria. FL-RhCMV-based vectors will thus be highly useful for both basic and translational aspects of CMV research.

## Materials and Methods

### Cells and viruses

Telomerized rhesus fibroblasts (TRFs) have been described before (63). Primary embryonal rhesus fibroblasts were generated at the ONPRC. Both cell lines were maintained in DMEM complete, Dulbecco’s modified Eagle’s medium (DMEM) with 10% fetal bovine serum and antibiotics (1× Pen/Strep, Gibco), and grown at 37°C in humidified air with 5% CO_2_. Rhesus retinal pigment epithelial (RPE) cells were a kind gift from Dr. Thomas Shenk (Princeton University, USA) and were propagated in a 1:1 mixture of DMEM and Ham’s F12 nutrient mixture with 5% FBS, 1 mM sodium pyruvate, and nonessential amino acids. Monkey kidney epithelial (MKE) (64) cells were maintained in DMEM-F-12 medium (DMEM-F12) (Invitrogen) supplemented with epithelial cell growth supplement (ScienCell), 1 mM sodium pyruvate, 25 mM HEPES, 100 U/ml penicillin, 100g/ml streptomycin, 2 mM L-glutamine (Invitrogen), and 2% fetal bovine serum/SuperSerum (Gemini Bio-Products). The RhCMV 68-1 BAC (22) has been characterized extensively. The BAC for RhCMV 68-1.2 (25) which was based on the 68-1 BAC was provided by Dr. Thomas Shenk (Princeton University, USA). Both 68-1 and 68-1.2 were derived via electroporation (250V, 950µF) of BAC-DNA into primary rhesus fibroblasts. Full cytopathic effect (CPE) was observed after 7-10 days and the supernatants were used to generate viral stocks. RhCMV UCD52 and RhCMV UCD59 have been continuously passaged on MKE cells to maintain their PRC and to minimize tissue culture adaptations. To generate enough viral DNA for a full genome analysis, monolayers of MKE at 90 – 100% confluence were inoculated with RhCMV (MOI: 0.01). Infections progressed to ∼90% CPE at which time supernatant (SN) and cells were collected and centrifuged at 6000 x g for 15 minutes at 4°C. SN was passed through a 0.45µm filter. SN was then centrifuged at 26,000 x g for 2 hours at 4°C. The SN was decanted and the virus pellet was resuspended and washed in ∼ 20ml of cold 1X PBS. Virus was pelleted by ultracentrifugation at 72,000 x g (Rotor SW41Ti at 21,000 rpm) for 2 hours at 4°C. This was repeated once more. SN was decanted and the remaining viral pellet was thoroughly resuspended in ∼1-2 ml of cold 1X PBS. Viral stock was stored in 50µl aliquots. Viral DNA was isolated from these viral stocks using the QIAamp DNA Mini Kit (Qiagen).

### BAC recombineering using en passant homologous recombination

Recombinant RhCMV clones were generated by *en passant* mutagenesis, as previously described for HCMV (65), and adapted by us for RhCMV (66). This technique allows the generation of “scarless” viral recombinants, i.e. without containing residual heterologous DNA sequences in the final constructs. The homologous recombination technique is based on amplifying an I-SceI homing endonuclease recognition site followed by an aminoglycoside 3-phosphotransferase gene conferring kanamycin resistance (KanR) with primers simultaneously introducing a homology region upstream and downstream of the selection marker into the intermediate BAC cloning product. As en passant recombinations are performed in the GS1783 *E-coli* strain that can be used to conditionally express the I-SceI homing endonuclease upon arabinose induction (65), expression of the endonuclease with simultaneous heat shock induction of the lambda (λ) phage derived Red recombination genes will lead to selective DNA double strand breaks with subsequent scarless deletion of the selection marker. The immunologically traceable markers used in the study, namely the SIVmac239 GAG protein as well as the *M.tuberculosis* Erdman strain derived Tb6Ag fusion protein, have been described before (14, 53). To introduce these genes into the FL-RhCMV backbone, we first introduced a homology region flanking an I-SceI site and a KanR selection marker into the selected inserts. We then amplified the transgenes by PCR and recombined the entire insert into the desired location in the FL-RhCMV BAC. The KanR cassette was subsequently removed scarlessly as described above. All recombinants were initially characterized by XmaI restriction digests and Sanger sequencing across the modified genomic locus. Lastly all vectors were fully analyzed by next generation sequencing to exclude off-target mutations and to confirm full accordance of the generated with the predicted full genome sequence.

### Isolation of the old world non-human primate CMV species from urine samples

Virus isolation was performed as previously described (39). Briefly, urine samples were obtained through collection from cage pans, by cystocentesis or following euthanasia. From samples collected at the Oregon National Primate Research Center (ONPRC) we isolated BaCMV 31282 from a male olive baboon (*Papio anubis*), BaCMV 34826 from a female hamadryas baboon (*Papio hamadryas*), CyCMV 31709 form a female cynomolgus macaques (*Macaca fascicularis)* of Cambodian origin, JaCMV 24655 from a male Japanese macaque (*Macaca fuscata*) and RhCMV 34844 from a male rhesus macaque (*Macaca mulatta*) of Indian origin. Additionally, we successfully isolated RhCMV KF03 from a cage pan collected urine sample from a male rhesus macaque (*Macaca mulatta*) of Indian origin housed at the Tulane National Primate Research Center (TNPRC). All urine samples were first clarified from solid contaminants by centrifugation at 2,000 x g for 10 minutes at 4°C and then filtered through a 0.45 μm filter (Millipore) to clear the urine of any bacterial or fungal contamination. Next, we spin-inoculated 0.5 ml – 2 ml of clarified urine onto primary rhesus fibroblasts in a 6 well plate at 700 x g for 30 minutes at 25°C. The cells were placed on a rocker for 2 hours at 37°C and, after removing the inoculum, washed once with PBS. The infected cells were cultured in DMEM plus 10% fetal bovine serum for 2-3 days, trypsinized and seeded in a T-175 cell culture flask. All samples were monitored weekly for CPE for up to six weeks or until plaque formation was visible. Every two weeks or after the appearance of plaques, cells were trypsinized and re-seeded to facilitate viral spread through the entire monolayer. Virus propagation was kept to an absolute minimum and viral stocks were prepared with the minimum number of passages required to be able to infect eight T-175 flasks for stock production (typically 1-3 passages).

### Isolation and purification of viral DNA for next generation sequencing (NGS)

The modified Hirt extraction (67) protocol used for the preparation of CMV viral DNA has been described (39). Briefly, supernatants from cells that were spin-inoculated with the original urine sample were collected at full CPE and used to infect three T-175 flasks of primary rhesus fibroblasts. After 7–10 days, the supernatant was harvested and clarified by centrifugation, first at 2,000 x g for 10 minutes at 4°C and subsequently at 7,500 x g for 15 minutes. Virus was pelleted through a sorbitol cushion (20% D-sorbitol, 50 mM Tris [pH 7.4], 1 mM MgCl_2_) by centrifugation at 64,000 x g for 1 hour at 4°C in a Beckman SW28 rotor. The pelleted virus was resuspended in 500μl 10.1 TE Buffer (10mM Tris, pH 8.0; 0.1mM EDTA, pH 8.0) and 500 μl 2x lysis buffer (20mM Tris-Cl, pH 8.0; 50mM EDTA, pH8.0; 200mM NaCl; 1.2% w/v SDS) was added. To digest the purified virion and to release the viral DNA, 250μg Proteinase K was added and the solution was incubated for 2h at 37°C. This was followed by two rounds of phenol/chloroform extractions and the viral DNA was precipitated overnight with absolute ethanol at −80°C. The DNA was pelleted for 20 minutes at 13,200 x g at 4°C, washed once with 70% ethanol, and subsequently resuspended in autoclaved double deionized water. DNA concentrations were determined using a ND-1000 Spectrophotometer (NanoDrop Technologies, Inc.).

### Generation of Next Generation Sequencing libraries and Next Generation sequencing

Illumina sequencing libraries were generated by first fragmenting the viral DNA using an S2 Sonicator and by subsequently converting the fragments into libraries using the standard TruSeq protocol. All libraries were quality controlled on a Bioanalyzer (Agilent) and library concentration was determined by real time PCR and SYBR Green fluorescence. Finally, the next generation sequencing was performed on either an iSeq- or a MiSeq Next-Generation Sequencing platform (Illumina). The libraries were multiplexed at equal concentrations and loaded into a reagent cartridge at 9 pM and single read sequencing was performed for 300 cycles with 6 additional cycles of index reads. The Geneious Prime software was used for all NGS data analysis. To minimize sequencing errors, the sequencing reads were trimmed of all regions exceeding the error probability limit of 0.1%. All reads with a total length of less than 50 bp after trimming were eliminated from further analysis to decrease the background due to unspecific alignment of reads during *de novo* and reference guided assemblies. All full genome sequences were first *de novo* assembled using the processed sequencing data, and subsequently all reads were aligned to the generated consensus sequence in a reference-guided assembly. All detected single nucleotide polymorphisms (SNPs) that showed a frequency of more than 50% in a location with a depth of at least 10% of the mean depth were manually curated. All nucleotide changes that were considered to be likely the results of actual changes opposed to sequencing errors or software alignment issues, were changed in the consensus sequences and the referenced guided assemblies were repeated until no SNP showed a frequency of 50% or more. The resulting final sequence was considered the representative consensus sequences of all clones contained in the primary viral isolates.

### Nucleotide sequences used and generated in this study

All full genome old world NHP CMV sequences generated in this study were submitted to GenBank. The accession numbers for the submitted isolates are as follows: BaCMV 31282 (MT157321),BaCMV 34826 (MT157322), CyCMV 31709 (MT157323), JaCMV 24655 (MT157324), RhCMV 34844 (MT157328), RhCMV KF03 (MT157329), RhCMV UCD52 (MT157330) and RhCMV UCD59 (MT157331). Furthermore, we submitted an updated annotation for the RhCMV 68-1 BAC (MT157325), a full annotation for the partially repaired RhCMV 68-1.2 BAC (MT157326) as well as a full annotation for the FL-RhCMV BAC described here (MT157327) which was based on 68-1 and 68-1.2. Genome annotations were fine-tuned utilizing these and other NHP CMV sequences that had been previously submitted to GenBank, either by us: CyCMV 31906 (KX689263), CyCMV 31907 (KX689264), CyCMV 31908 (KX689265), CyCMV 31909 (KX689266), RhCMV 180.92 (DQ120516, AAZ80589), RhCMV 19262 (KX689267), RhCMV 19936 (KX689268) and RhCMV 24514 (KX689269) or by others: CyCMV Ottawa (JN227533, AEQ32165), CyCMV Mauritius (KP796148, AKT72642), SCMV Colburn (FJ483969, AEV80601), SCMV GR2715 (FJ483968, AEV80365), SCMV Stealth virus 1 strain ATCC VR-2343 (U27469, U27238, U27770, U27627, U27883, U27471), BaCMV OCOM4-37 (AC090446), BaCMV OCOM4-52 (KR351281, AKG51610.1), DrCMV OCOM6-2 (KR297253, AKI29779). Lastly, we also used full genome sequences of CMV species from outside the old world NHPs in our phylogenetic analysis to classify the evolutionary relationship of FL-RhCMV to other CMVs. These additional sequences include: MCMV Smith (GU305914, P27172), RCMV Maastricht (NC_002512), RCMV England (JX867617), RCMV Berlin (KP202868), GPCMV 22122 (KC503762, AGE11533), AoHV-1 S34E (FJ483970, AEV80760), SaHV-4 SqSHV (FJ483967, AEV80915), CCMV Heberling (AF480884, AAM00704), and HCMV TR3 (MN075802).

### Phylogenetic analysis of isolated NHP CMV species

Full nucleotide sequences of rodent and primate CMV genomes were aligned using ClustalW2 (68). The multiple sequence alignment was imported into Geneious Prime and a Neighbor-joining phylogenetic tree was build using the Geneious Tree Builder application and selecting the Jukes-Cantor genetic distance model using the MCMV Smith strain as an outgroup. The validity of the tree topology obtained was tested by using bootstrap analysis with 100 resamplings from the aligned sequences, followed by distance matrix calculations and calculation of the most probable consensus tree with a support threshold of 50%.

### Conditional expression of the Rh13.1 (RL13) encoded mRNA using the Tet-Off system

TRF were engineered to express the tetracycline-repressor (tetR) as previously described (69, 70). Briefly, a retrovirus was generated from vector pMXS-IP expressing the tetR ORF. Retronectin was then used to transduce TRFs with high efficiency, before selection in puromycin (1μg/ml). Successful expression of the tetR was validated using replication deficient recombinant adenovirus vectors expressing GFP from a tetO-regulated promoter (70).

To create a vectors conditionally expressing Rh13.1, we inserted dual tetO sequences 131bp upstream of its start codon. Next, eGFP was inserted as a marker of infection. eGFP was linked to Rh60/61 (homologous to HCMV UL36) via a P2A linker. We have previously shown for HCMV that this provides early expression of GFP, without affecting UL36 function (6). As a control, we created a virus in which we deleted the entire Rh13.1 ORF. All vectors were created using *en passant* mutagenesis.

To analyze the effects of Rh13.1 on plaque formation, BAC DNA was transfected into TRFs or tetR-expressing TRFs using an Amaxa Nucleofector (basic fibroblast kit, program T-16). The formation of plaques was then monitored by imaging for eGFP fluorescence at various timepoints, using a Zeiss Axio Observer Z1.

### Rh13.1 (RL13) mRNA regulation using riboswitches

To generate a FL-RhCMV vectors carrying riboswitches, we inserted the published Tc45 aptazyme sequence 19bp upstream and the Tc40 aptazyme sequence 17bp downstream of Rh13.1 (45) into the BAC by incorporating them into the homologous recombination primers. The entire 122bp sequences were introduced into the 5’ and the 3’ flanking regions of Rh13.1 in two independent recombination steps. The resulting BAC construct was analyzed by XmaI endonuclease restriction digest, Sanger sequencing and next generation sequencing.

To reconstitute the virus, we transfected BAC DNA into primary rhesus fibroblast using Lipofectamine 3000 (ThermoFischer) in the presence of 100 µM tetracycline which was replenished every other day. For comparison, we reconstituted virus in the absence of tetracycline. After minimal passaging in the presence or absence of tetracycline virus stocks were generated, viral DNA was isolated and NGS was performed.

### Multi-step growth curves of RhCMV on primary rhesus fibroblasts

Primary rhesus fibroblast were seeded in 24 well plates and infected in duplicate with RhCMV constructs at an MOI of 0.01. The inoculum was removed after 2h and 1ml DMEM complete was added. Supernatants were collected every third day for 24 days, cells and cell debris were removed by centrifugation for 2 min at 13,000 rpm and the samples were stored at −80°C. Viral titers were determined using a fifty-percent tissue culture infective dose (TCID50) assay in two technical repeats. Final titers were calculated using the arithmetic mean of the technical and biological repeats.

### RhCMV entry assays into primary rhesus fibroblast and rhesus retinal pigment epithelial cell

Stocks for RhCMV strains 68-1, 68-1.2 and FL-RhCMV were generated and viral titers were determined by TCID50. Infection levels in fibroblasts were experimentally equalized across stocks. Primary rhesus fibroblast were infected at an MOI of 0.3 and RPEs at an MOI of 10 as these MOIs were experimentally determined to result in optimal infection levels. The cells were fixed at 48 hpi and the overall percentage of RhCMV positive cells were determined by flow cytometry using a RhCMV specific antibody (52). To be able to compare infection levels between the two cell types, we set the mean infection level in fibroblasts determined in triplicate repeats to 100% and expressed the mean infection level of triplicate repeats in RPEs in relation to the infection level in fibroblasts.

### Quantitative PCR (qPCR) analysis to assess mRNA expression levels

Primary rhesus fibroblasts were seeded in 6-well plates and infected either with FL-, 68-1, or 68-1.2 RhCMV at a MOI of 1. Total RNA was then isolated at 48 hours post infection (hpi). Uninfected rhesus fibroblasts were used as a negative control. After cDNA synthesis, the quantitative PCR (q-PCR) assay was performed using primers and probes specific to each gene of interest. Rh159_forward_primer: 5’ TCAGAAATGAAGGGCAATTGTG 3’. Rh159_reverse_primer: 5’ GCGAGCTGGCGACGTT 3’. Rh159_probe: 6FAMTATCACTCGGCTATTATCMGBNFQ. Rh137_forward_primer: 5’ GGCGCAACATACTACCCAGAA 3’. Rh137_reverse_primer: 5’ GTAGCCATCCCCATCTTCCA 3’. Rh137_probe: 6FAMCACAACTAACTCTGGCCTTMGBNFQ. GAPDH_forward_primer: 5’ TTCAACAGCGACACCCACTCT 3’. GAPDH_reverse_primer: 5’ GTGGTCGTTGAGGGCAATG 3’. GAPDH_probe: 6FAMCCACCTTCGACGCTGGMGBNFQ. The mRNA copy numbers for Rh159 (UL148) and Rh137 (UL99) were calculated and graphed as relative mRNA copy numbers normalized to the housekeeping gene (GAPDH).

### Ethics statement

All RMs housed at the Oregon National Primate Research Center (ONPRC) and the Tulane National Primate Research Center (TNPRC) were handled in accordance with good animal practice, as defined by relevant national and/or local animal welfare bodies. The RMs were housed in Animal Biosafety level (ABSL)-2 rooms. The rooms had autonomously controlled temperature, humidity, and lighting. Study RM were both pair and single cage-housed. Regardless of their pairing, all animals had visual, auditory and olfactory contact with other animals within the room in which they were housed. Single cage-housed RMs received an enhanced enrichment plan and were overseen by nonhuman primate behavior specialists. Animals were only paired with one another if they were from the same vaccination group. RMs were fed commercially prepared primate chow twice daily and received supplemental fresh fruit or vegetables daily. Fresh, potable water was provided via automatic water systems. The use of nonhuman primates was approved by the ONPRC and the TNPCR Institutional Animal Care and Use Committees (IACUC). Both institutions are Category I facilities and they are fully accredited by the Assessment and Accreditation of Laboratory Animal Care International and have an approved Assurance (#A3304-01) for the care and use of animals on file with the NIH Office for Protection from Research Risks. The IACUCs adhere to national guidelines established in the Animal Welfare Act (7 U.S.C. Sections 2131–2159) and the Guide for the Care and Use of Laboratory Animals (8th Edition) as mandated by the U.S. Public Health Service Policy. For BAL procedures, monkeys were restrained and sedated by intramuscular injection of ketamine (∼7 mg/kg) with dexmedetomidine (∼15 μg/kg). Following the procedure (and blood collection), atipamezole (5 mg/mL; the dose volume was matched to that of dexmedetomidine) was administered by intramuscular injection to reverse the effects of the dexmedetomine sedation. To prepare RMs for blood collection alone, monkeys were administered ketamine only as described above. Monkeys were bled by venipuncture (from the femoral or saphenous veins) and blood was collected using Vacutainers. Monkeys were humanely euthanized by the veterinary staff at ONPRC and TNPCR in accordance with end point policies. Euthanasia was conducted under anesthesia with ketamine, followed by overdose with sodium pentobarbital. This method is consistent with the recommendation of the American Veterinary Medical Association.

### Intravenous (i.v.) inoculation of CMV naïve RM with FL-RhCMV

Three immunocompetent male RhCMV-seronegative Indian-origin rhesus macaques from the expanded SPF colony at the TNPRC were intravenously inoculated with 1.79×10^6^ pfu of Rh13.1-intact FL-RhCMV/Rh13.1/apt. Blood, saliva samples collected by oral sterile PBS wash, and urine from cage pans were collected at biweekly to weekly intervals until necropsy 9 weeks post RhCMV inoculation. Plasma samples were shipped to UC Davis on dry ice and stored at −80°C until use. For DNA isolation, they were quickly thawed for up to 1 minute in a 37°C dry bath. Each plasma sample was vortexed for 15 seconds and then 350µl of the plasma was transferred to a sterile, 2.0 ml polypropylene screw cap tube (Sarstedt, 72.693.005). Plasma tubes were loaded into the QIAsymphony SP (Qiagen, Germantown, MD) and purified DNA was obtained using the DNA Blood 200µl protocol and the QIAsymphony DNA Mini kit reagent cartridge (Qiagen, 931236). Quantitative PCR was performed on the plasma derived DNA using the 7900HT Real-Time PCR System (Applied Biosystems, Inc. and Thermo Fisher, Grand Island, NY), and a previously published protocol for qPCR using primers specific to RhCMV gB (71). Results were analyzed using the Sequence Detection Systems software (vs. 2.4). Fluorescence intensity in each well was measured, and a result was considered positive when the intensity exceeded 10 times the baseline fluorescence. The limit of sensitivity was reproducibly between 1 and 10 copies of template DNA. Plasma DNA viral loads were calculated as copy number per milliliter of plasma. Plasma viral load data in the FL-RhCMV inoculated RM was compared with historical control pregnant female macaques inoculated with RhCMV strains UCD52, UCD59, and 180.92 as previously reported (11). RhCMV DNA copy numbers in these animals were quantitated by real time PCR using primers and probe targeting exon 1 of the immediate early gene of RhCMV using previously published methodology (72).

### Nested real-time PCR

To examine the differences in dissemination between FL-RhCMVΔRh13.1/TB6Ag and FL-RhCMVΔRh13.1/TB6AgΔRh157.4-5ΔRh158-161, we infected three CMV naïve RM per vector s.c. with 10^7^ PFU. At 14 days post infection, we took the animals to necropsy and harvested tissues from which the DNA was isolated by the ONPRC Molecular Virology Support Core (MVSC) using the FastPrep (MP Biomedicals) in 1 ml TriReagent (Molecular Research Center Inc.) for tissue samples under 100 mg. Additionally, 100 μl bromochloropropane (MRC Inc.) was added to each homogenized tissue sample to enhance phase separation. 0.5 ml DNA back-extraction buffer (4 M guanidine thiocyanate, 50 mM sodium citrate, and 1 M Tris) was added to the organic phase and interphase materials, which was then mixed by vortexing. The samples were centrifuged at 14,000 g for 15 minutes, and the aqueous phase was transferred to a clean microfuge tube containing 240 μg glycogen and 0.4 ml isopropanol and centrifuged for 15 minutes at 14,000 g. The DNA precipitate was washed twice with 70% ethanol and resuspended in 100 to 500 μl double deionized water. Nested real-time PCR was performed with primer and probe sets for the inserted *Mtb* Tb6Ag transgene (first round: for-CAGCCGCTACAGATGGAGAG and rev-CGCGCTAGGAGCAAATTCAC; second round: for-CAGCCGCTACAGATGGAGAG, rev-CGCGCTAGGAGCAAATTCAC, and probe-TGGCGGCTTGCAAT-FAM). For each DNA sample, 10 individual replicates (5 μg each) were amplified by first-round PCR synthesis (12 cycles of 95°C for 30 seconds and 60°C for 1 minute) using Platinum Taq in 50 μl reactions. Then, 5 μl of each replicate was analyzed by nested quantitative PCR (45 cycles of 95°C for 15 seconds and 60°C for 1 minute) using Fast Advanced Master Mix (ABI Life Technologies) in an ABI StepOnePlus Real-Time PCR machine. The results for all 10 replicates were analyzed by Poisson distribution and expressed as copies per cell equivalents (73).

### Histopathology and in situ hybridization (ISH)

RNAscope was performed on formaldehyde fixed, paraffin-embedded tissue sections (5μm) according to our previously published protocol (74) with the following minor modifications: heat-induced epitope retrieval was performed by boiling slides (95-98°C) in 1x target retrieval (ACD; Cat. No. 322000) for 30 min., followed by incubation at 40°C with a 1:10 dilution of protease III (ACD; Cat. No. 322337;) in 1x PBS for 20 min. Slides were incubated with the target probe RhCMV (ACD; Cat. No. 435291) for 2 hours at 40°C and amplification was performed with RNAscope 2.5 HD Brown Detection kits (ACD; Cat. No. 322310) according to manufacturer’s instructions, with 0.5X wash buffer (310091; ACD) used between steps, and developed with Alexa-fluor647 conjugated tyramide. The RhCMV probe consisted of 50zz pairs targeting the RhCMV rh38 (13zz pairs), rh39 (10zz pairs all shared with Rh39) and rh55 (37zz pairs) ORFs originally annotated in RhCMV 68-1 (55). These ORFs correspond to Rh38.1 (UL22A) and Rh55 (UL33) in the here presented annotation of the RhCMV genome. To remove/inactivate the *in situ* amplification tree/HRP complexes, slides were microwaved at full power for 1 minute, followed immediately for 15 minutes at 20% power in 0.1% citraconic anhydride with 0.05% Tween-20 buffer. Slides were cooled for 15 minutes in the buffer, then rinsed in ddH2O. Slides were subsequently stained for CD34 (Sigma, Cat. No. HPA036723), at a 1:200 dilution in antibody diluent (1x TBS containing 0.25% casein and 0.05% Tween-20) overnight at 4°C. Detection was performed using an anti-rabbit polymer HRP conjugated system (GBI Labs; Cat. No. D39-110), and developed with Alexa-fluor488 conjugated tyramide at a 1:500 dilution for 10 minutes. To remove the antibody/HRP complexes, a second round of microwaving was performed as described above. Slides were subsequently stained for myeloid lineage cells using a combination of mouse anti-CD68 (Biocare Medical; Cat. No. CM033C; clone KP1) and mouse anti-CD163 (ThermoFisher; Cat. No. MA5-1145B; clone 10D6), at a 1:400 dilution (each) in antibody diluent for one hour at room temperature. Detection was performed using an anti-mouse polymer HRP conjugated system (GBI Labs; Cat. No. D37-110), and developed with Alexa-fluor350 conjugated tyramide at a 1:50 dilution for 15 minutes. A third round of microwaving was performed to remove the antibody/HRP complexes as described above. Slides were subsequently stained for mesenchymal lineage cells using a mouse anti-vimentin (Sigma; Cat. No. HPA001762), at a 1:1000 dilution in antibody diluent overnight at 4°C. Detection was performed using an anti-mouse polymer HRP conjugated system (GBI Labs; Cat. No. D12-110), and developed with Alexa-fluor594 conjugated tyramide at a 1:500 dilution for 10 minutes. To ensure that that HRP inactivation and antibody stripping was complete, matched slides that had gone through the previous staining steps did not receive the subsequent primary antibody, but were developed with all slides from that round. In each case we did not see staining with Alexa-fluor 488, Alexa-fluor 350 or Alexa-fluor 594 tyramide, indicating that the microwave step completely removed/inactivated the *in situ* amplification tree/HRP complexes and removed all previous antibody/HRP complexes. Slides were coverslipped using Prolong Gold antifade mounting media (ThermoFisher; Cat. No. P36930), scanned using a Zeiss AxioScan Z1, and analyzed using Halo software (v2.3.2089.52; Indica Labs). Cell counts were quantified using the FISH Multiplex RNA v2.1 module and RhCMV RNA percent area quantification was performed using the Area Quantification FL v1.2 module. To calculate viral copy number, we used the HALO analysis program to analyze the average size (area) and fluorescent intensity of more than 10 individual virions within the spleen, which was used that to calculate a minimum estimate of RhCMV RNA copy numbers in the infected cells using the FISH Multiplex RNA v2.1 module. Statistical analysis was performed with GraphPad Prism v.7.04. Data are mean +/- S.E.M. as indicated.

### Intracellular cytokine staining (ICS)

Our intracellular cytokine staining (ICS) protocol to examine immune cells isolated from RM tissues has been described previously (14, 75). Briefly, we isolated mononuclear cells from RM tissues collected at necropsy and measured specific CD4^+^ and CD8^+^ T cell responses by flow cytometric ICS. For this purpose, the isolated cells were incubated with mixtures of consecutive 15mer peptides (overlapping by 11AA) of the *Mtb* Tb6Ag in the presence of the costimulatory molecules CD28 and CD49d (BD Biosciences) for 1 hour, followed by addition of brefeldin A (Sigma-Aldrich) for an additional 8 hours. As a background control we used cells co-stimulated without the peptide pool. Following incubation, cells were stored at 4°C until staining with combinations of fluorochrome-conjugated monoclonal antibodies including: anti-CD3 (SP34-2: Pacific Blue; BD Biosciences), anti-CD4 (L200: BV510; Biolegend), anti-CD8α (SK1: TruRed; eBioscience), anti-TNF-α (MAB11: PE; Life Tech), anti-IFN-γ (B27: APC; Biolegend) and anti-CD69 (FN50: PE/Dazzle 594; Biolegend), anti-Ki67 (B56: FITC; BD Biosciences). Data were collected on an LSR-II flow cytometer (BD Biosciences). FlowJo software (Tree Star) was used for data analysis. In all analyses, gating on the small lymphocyte population was followed by the separation of the CD3^+^ T cell subset and progressive gating on CD4^+^ and CD8^+^ T cell subsets. Antigen-responding cells in both CD4^+^ and CD8^+^ T-cell populations were determined by their intracellular expression of CD69 and either or both of the cytokines interferon gamma (IFN-γ) and tumor necrosis factor alpha (TNF-α). Final response frequencies shown have been background subtracted and memory corrected as previously described (76).

### RNA-seq Library Preparation, Sequencing and Data Analysis

Total RNA was isolated from *in vitro* cell cultures and *in vivo* tissues using the Trizol method. RNA next generational sequencing (NGS) was performed on polyA-fractionated RNA utilizing the TruSeq Stranded mRNA library prep kit (Illumina). Library was validated using Agilent DNA 1000 kit on bioanalyzer according to manufacturer’s protocol. RNA libraries were sequenced by the OHSU Massively Parallel Sequencing Shared Resource Core using their Illumina HiSeq-2500 using single-end 100 bp reads. Due to low fraction of viral reads relative to host, the libraries from parotid and submandibular glands were sequenced again to increase read depth. Sequence data were quality trimmed with Trimmomatic (77), and aligned to a custom reference genome comprised of the latest rhesus macaque genome build (MMul10; assembly ID GCA_003339765.3) and the RhCMV68-1/Tb genome using the STAR aligner (78). For coverage analyses, GATK DepthOfCoverage (79) was used to produce a table of raw counts per base. Coverage across the RhCMV genome was visualized in R (3.6.1) for the three in vitro dataset (8, 24, and 72 hour post infection) using Gviz (1.3) (80). Base-pair level coverage data was smoothed using the ksmooth function of base R, a kernel-based regression smoothing technique, with the bandwidth parameter set to 500. Feature counting was performed using Subread featureCounts version 1.6.0 (81), using a GTF produced by concatenating Rhesus macaque Ensembl reference genes (build 98) with the ORFs annotated from RhCMV-68.1/Tb. For subread, the following options were used: fracOverlap of 0.1, minOverlap of 20, using both --largestOverlap, and --primary. To account for gene length differences that can bias the transcript counts, we normalized across the genes, for all samples by the gene length (computed as the total exon length from start to end positions). Finally, for an equivalent comparison across the *in vitro* and *in vivo* samples, used for the principal component analysis (PCA) as well as the combined heatmap (Fig.8A and B), we corrected for cross-sample variation by quantile normalization of the gene expression matrix. PCA was performed using the base R prcomp function on the normalized gene expression matrix as described in the data analysis section. The variance across the components was used to order and select for top components of interest.

## Author Contributions

Conceptualization: HT, EM, BNB, LJP, DNS, KF, DM

Data Curation: HT, EM, SGH, AK, BNB, LJP, DNS, DM

Formal Analysis: HT, EM, MN, JS, PTE, SGH, BNB, DM

Funding Acquisition: LJP, KF

Investigation: HT, EM, CK, LSU, MRM, MJM, AB, MN, TW, EAS, LMS, DR, CMH, KAJ, ANS, ABV, YY, KS, RJS, SGH, BNB, DM

Methodology: EM, BNB, LJP, KF, DM

Project Administration: RJS, SGH, AK, DNS, DM

Resources: JS, MKA, JDE, SGH, AK, PAB, BNB, LJP

Software: EM, BNB

Supervision: SGH, DNS, DM

Validation: HT, EM, BNB, DM

Visualization: HT, EM, MJM, MN, SGH, BNB, DM

Writing – Original Draft Preparation: KF, DM

Writing – Review & Editing: HT, EM, MJM, RJS, JDE, SGH, AK, PAB, BNB, LJP, DNS, KF, DM

## Acknowledgements

We would like to thank Dr. Thomas Shenk (Princeton University) for providing the RhCMV 68-1.2 BAC and the rhesus retinal pigment epithelium (RPE) cells. We would also like to thank the faculty and staff of the Departments of Veterinary Medicine and Collaborative Research at TNPRC for the excellent animal care and collection of research specimens. We are grateful to the OHSU molecular virology support core for generating RhCMV stocks and for assisting in the processing of tissue samples, to Dr. Robert Searles and the OHSU Massively Parallel Sequencing Shared Resource (MPSSR) for generating the NGS libraries and to Yibing Jia and the Molecular and Cellular Biology Core (MCB Core) at the ONPRC for analyzing the samples on their MiSeq next generation sequencer. Lastly, we would like to acknowledge the invaluable technical and scientific support provided by Katinka Vigh-Conrad, Larry Wilhelm, Shoko Hagen, Andrew Sylwester, Kyle Taylor, Junwei Gao, Jennie Womack, Nessy John and Hillary Cleveland-Rubeor that made this study possible. This work was supported by the National Institute of Allergy and Infectious Diseases (NIAID) (P01 AI129859U42, R01 AI095113, P01 AI094417, R37 AI054292, R01 AI059457, OD023038), the National Institutes of Health Office of the Director (U42OD010426, P51OD011092, P51OD011104), the Eunice Kennedy Shriver National Institute of Child Health & Human Development (NICHD) (4DP2HD075699) and the Bill & Melinda Gates Foundation (OPP1033121, OPP1108533, OPP1152430).

---

**Figure S1: Genome organization of NHP CMVs**

The genome of HCMVs and the closely related CCMV comprise two unique coding regions (U_L_ and U_S_) that are separated by an internal repeat region and flanked by terminal repeats. The repeat regions consist of the three repeated sequence units a, b and c that form overlapping inverted repeats in the form *ab-U_L_-b’a’c-U_S_-c’a*. The HCMV genome can re-arrange to four different isomers varying in the relative orientation of the U_L_ and U_S_ regions to one another (97). Intriguingly, while the U_L_ and U_S_ regions can still be identified in old world NHP CMVs, the repeat organization is vastly different. The terminal direct repeats in these species are short while the internal repeats are completely missing resulting in a single isomer that has been fixed during evolution. All Asian NHP CMVs and the African green monkey (Simian) CMV (SCMV) occur in the same isomeric form whereas the U_S_ region appears in the opposite orientation to the U_L_ region in the closely related BaCMV and DrCMV. New world (NW) CMVs retained a genome organization with terminal and internal repeats similar HCMV, but the repeats are organized as non-overlapping inverted repeats flanking the U_L_ and U_S_ regions, allowing for isomerization.

**Figure S2: The RL11 gene family of NHP CMVs**

A) Phylogenetic tree of RL11 family genes from representatives of each NHP CMV species. B) ORF structure of the RL11 family genes in each NHP CMV species. Each gene is color-coded using the same colors as in A) showing the presence/absence of each ORF in a given NHP CMV species.

**Figure S3: The UL83 (pp65) gene family of NHP CMVs**

A) Phylogenetic tree based on the protein sequences of NHP CMV genes homologous to HCMV UL83 encoding the major tegument protein pp65 from representatives of each NHP CMV species. B) ORF structure of the UL83 family genes in each NHP CMV species. Each gene is color-coded using the same colors as in A) showing the presence/absence of each ORF in a given NHP CMV species.

**Figure S4: The UL146/UL147 gene family of NHP CMVs**

A) Phylogenetic tree based on the protein sequences of NHP CMV genes homologous to HCMV chemokine-like genes UL146 and UL147 from representatives of each NHP CMV species. B) ORF structure of the UL146/147 family genes in each NHP CMV species. Each gene is color-coded using the same colors as in A) showing the presence/absence of each ORF in a given NHP CMV species.

**Figure S5: The Rh166 gene family of NHP CMVs**

A) Phylogenetic tree based on the protein sequences of Rh166 family genes from representatives of each NHP CMV species. B) ORF structure of the Rh166 family genes in each NHP CMV species. Each gene is color-coded using the same colors as in A) showing the presence/absence of each ORF in a given NHP CMV species.

**Figure S6: The US12 gene family of NHP CMVs**

A) Phylogenetic tree based on the protein sequences of NHP CMV genes homologous to the HCMV US12 family encoding seven transmembrane proteins from representatives of each NHP CMV species. B) ORF structure of the US12 family genes in each NHP CMV species. Each gene is color-coded using the same colors as in A) showing the presence/absence of each ORF in a given NHP CMV species.

**Figure S7: The US28 gene family of NHP CMVs**

A) Phylogenetic tree based on the protein sequences of NHP CMV genes homologous to HCMV US28 encoding G-protein coupled receptors from representatives of each NHP CMV species. B) ORF structure of the US28 family genes in each NHP CMV species. Each gene is color-coded using the same colors as in A) showing the presence/absence of each ORF in a given NHP CMV species.

**Figure S8: Exploratory analysis to assess equivalency across RNA-seq data.**

To account for equivalency across the RNA-seq samples, A) background expression was assessed using house-keeping genes (“ACTG1”, “RPS18”, “MRPL18”, “TOMM5”, “YTHDF1”, “TPT1”, “RPS27”) (98) which did not identify a specific trend across samples. Next, B) host (RM) library size was assessed across samples which showed higher overall transcript levels that were tissue specific to the salivary glands. However, this difference did not correlate with the viral gene expression levels. Finally, C) we found that all three in vitro samples had a higher number of total viral reads than tissue biopsies.

**Figure S9: Sequence coverage of FL-RhCMVΔRh13.1/TB6Ag in cultured fibroblasts**

Sequence coverage of FL-RhCMVΔRh13.1/TB6Ag in cultured fibroblasts, infected *in vitro* and sampled at three timepoints post-infection (8, 24, 72 hours). Coverage per base is plotted with a log10 scale. Colors: blue = 8hpi, magenta = 24hpi, green = 72hpi.

**Table S1: List of all RhCMV ORFs affected by changes made to the previously published genome annotation using comparative genomics.**

**Table S2: Absolute number of RNAseq reads aligning to annotated RhCMV ORFs for all *in vitro* and *in vivo* samples.**

